# Age-dependent differential iron deficiency responses of rosette leaves during reproductive stages in *Arabidopsis thaliana*

**DOI:** 10.1101/2024.11.28.625839

**Authors:** Mary Ngigi, Mather Khan, Ricarda Remus, Shishir K Gupta, Petra Bauer

## Abstract

Iron (Fe) is essential for plant development throughout the life cycle. Rosette leaves are responsive to Fe supply in *Arabidopsis thaliana*. Little is known about the dynamics of Fe deficiency (-Fe) responses of rosette leaves during the reproductive stages. We studied the dynamics of Fe-dependent responses at four consecutive reproductive stages (rosette, bolting, flowering, mature silique stages, hereby named RS, BS, FS, MS). We examined the growth of rosette leaves, elemental contents and gene expression patterns of Fe homeostasis genes belonging to differently regulated groups. We determined individual leaf sizes during seven days of +Fe and -Fe treatment at the RS. Young leaves responded to -Fe with growth inhibition and yellowing. Old and young leaves differed in gene expression patterns and elemental contents. Differences were noted between the early and late reproductive stages (primarily RS and BS versus MS) and correlations between ionomic contents and gene expression were detected. All leaves had induced Fe recycling genes at -Fe.

Our findings highlight a developmental stage-dependent modulation of +Fe and -Fe responses in leaves. We discuss possible leaf signaling mechanisms accounting for the distinct responses between old and young leaves. This insight is informative to strengthen our understanding on plant iron management.

**Highlight/ One-sentence summary:** This study explores reproductive stage-dependent responses of rosette leaves to iron availability, revealing distinct growth, elemental contents, and gene expression patterns between young and old leaves and stages.

## Introduction

Iron (Fe) is a vital micronutrient for all organisms. In plants, Fe is central for the many fundamental Fe-requiring metabolic processes like photosynthesis, nutrient assimilation, chlorophyll synthesis, and oxidative stress defense (Briat et al., 2007; Jeong & Guerinot, 2009). Fe is often not bioavailable in soil and limiting during plant growth, requiring acclimation of plant Fe physiology. The complexities of iron homeostasis and networks of Fe-related cell physiological functions can be unraveled by exposing plants to Fe deficiency (-Fe) and recording growth data, ionomics profiles, and Fe-related gene expression patterns. Understanding Fe-regulatory pathways paves the way for innovations in translational plant biotechnology, such as to generate biofortified Fe-rich crops for food nutritional security that are suited for sustainable agroecological practices.

*Arabidopsis thaliana* has greatly served as a model species to generate knowledge about the regulation and genetic networks that steer the ways how Fe is acquired and allocated in plants (Sági-Kazár et al., 2022). Yet, there are some gaps in our understanding. Many studies focused on investigating the roles of Fe-regulatory genes, particularly in young seedlings and during the vegetative growth phase (Khan et al., 2018; Mai et al., 2016; Muhammad et al., 2022; Nguyen et al., 2022; Pu & Liang, 2023). The few studies, that examined Fe management during reproduction, concluded that *de novo* Fe uptake and Fe recycling are important to compensate for a lack of Fe, and that rosette leaves may act as Fe reserves that possibly supply Fe to newly developing organs through remobilization mechanisms (Pottier et al., 2019; Schuler et al., 2012; Waters & Grusak, 2008). Yet, to our knowledge, a detailed and systematic study that investigates how different rosette leaves of Arabidopsis respond to -Fe cues during the early and late reproductive stages is missing.

During the reproductive stages, the rosette leaves can be classified into old and young leaves, distinguishing them based on their age whereby first true leaves are fully matured at the onset of reproduction, while the later developed leaves continuously grow and expand until late reproductive stages. Old and young leaf types differ in morphology and size, and hints are available that they also show variation in the regulation of Fe homeostasis under changing nutrient conditions (Nguyen et al., 2022; Schuler et al., 2012; Swartz et al., 2023). Better understanding how different types of leaves manage Fe and Fe-related gene expression will provide new insights into leaf Fe signaling in Arabidopsis. Numerous mutants and tools are available in *A. thaliana*, that can be leveraged for further studies to better understand the genetic mechanisms directing Fe physiology during reproduction.

Gene co-expression analysis is a powerful tool to decipher Fe-regulatory networks, cell functions, and pathways for Fe management at Fe sufficiency (+Fe) and Fe deficiency (-Fe) conditions (Ivanov et al., 2012; Schwarz & Bauer, 2020). Co-expression indicates that different genes interact so that their collective expression leads to specific cellular and physiological outcomes being controlled by similar stimuli such as +Fe or -Fe. Of particular interest, the similar expression patterns of co-expressed -Fe-regulated genes across different conditions, time points, or tissues, indicate that they are most likely regulated by the same group of transcription factors and involved in related Fe regulatory processes (Schwarz & Bauer, 2020; Spielmann et al., 2023). Analyzing -Fe-induced co-expressed genes has helped to understand the complexities of Fe-regulatory systems. The Fe deficiency-induced co-expressed genes are divided into sub-categories of co-expression and co-regulation networks, that reflect the division into different Fe-related functions (Ivanov et al., 2012; Schwarz & Bauer, 2020). One subset of genes or co-expression cluster induced in response to -Fe in seedling roots serves to mobilize and acquire *de novo* Fe from soil, a process relevant in the root epidermis. This subset is also referred to as “FIT-dependent” since its up-regulation requires the presence of a transcription factor called FIT (FER-LIKE FE DEFICIENCY-INDUCED TRANSCRIPTION FACTOR). A second subset is often termed as “FIT-independent” since the expression of the genes in this cluster is particularly high when FIT function is lost leading to the inability of acquiring *de novo* Fe hence induction of -Fe signals in the plant. This subset of co-expressed genes is active in both, roots and shoots, and functions to mobilize and allocate internal Fe from root to shoot or within the shoot and this subset is primarily expressed in the root epidermis, root stele and in leaf vasculature (Long et al., 2010; Selote et al., 2015; Stacey et al., 2008; Wang et al., 2007). This co-expression cluster encodes proteins that recycle Fe from vacuolar stores, control intercellular Fe redistribution through phloem-based Fe-nicotianamine chelates, unloading towards Fe sinks (Bastow et al., 2018; Lanquar et al., 2005; Schuler et al., 2012), and sequester nicotianamine in root vacuoles for the management of metal ions (Haydon et al., 2012; Haydon & Cobbett, 2007). The concept of distinguishing co-expressed “FIT-dependent” and “FIT-independent” genes has been applied to classify the hierarchy of Fe responses and decipher signaling and regulatory events (Ivanov et al., 2012; Lichtblau et al., 2022; Muhammad et al., 2022; Schwarz & Bauer, 2020).

We hereby set out to investigate and correlate the -Fe responses of old and young rosette leaves in spatial- and time-resolved manner across four reproductive stages. We recorded growth dynamics, Fe-related gene expression changes and metal ion contents under +Fe and -Fe exposure. We hypothesized that reproductive transition profoundly impacts Fe homeostasis, and we predicted an increase in -Fe sensitivity in old and young leaves from early to late reproductive stages. However, this systematic study uncovered unexpected dynamics on reproductive and stage-specific Fe homeostasis regulation under changing Fe supply with distinct patterns in old and young leaves.

## Materials and Methods

### Plant materials and growth conditions

*Arabidopsis thaliana* ecotype Columbia (Col-0) seeds were surface-sterilized, stratified for 48 hours at 4°C in the dark and grown vertically on half-strength Hoagland agar plates (Lichtblau et al., 2022) in a growth chamber (CLF Plant Climatics, Wertingen, Germany) at 16 h/8 h photoperiod with 100 μmol m^−2^ s^−1^ light intensity, 22°C day/20°C night and 50 to 60 % relative humidity. 12-day-old seedlings were transferred into a hydroponic setup as previously described by (Nguyen et al., 2016) using a quarter-strength Hoagland hydroponic medium containing macronutrients (0.375 mM MgSO_4_, 0.75 mM Ca (NO_3_)_2_, 0.25 mM KH_2_PO_4_, 0.625 mM KNO_3_) and micronutrients (5 μM MnSO_4_, 1 μM ZnSO_4_, 0.75 μM CuSO_4_, 0.0375 μM (NH_4_)_6_Mo_7_O_24_, 25 μM KCl, and 25 μM H_3_BO_3_). 25 µM FeNaEDTA was added to the nutrient medium as sufficient Fe (+Fe). The medium was replaced every three days with freshly prepared hydroponic solution aerated using air pumps. Plants were grown up to four reproductive stages as previously described in (Boyes et al., 2001), hereby named the rosette stage (RS, early reproductive stage with <1cm inflorescence stem and closed inflorescence bud, 24-day-old), bolting stage (BS, inflorescence stem 2-4cm with closed inflorescence bud advanced inflorescence with beginning internode elongation, 32-day-old), flowering stage (FS, main inflorescence stem >5cm with 3-5 open floral buds, 38-day-old) and mature silique stage (MS, main inflorescence stem >8cm with 6-7 mature yellowing siliques, 46-day-old) (Figure 1A, Figure S1A, B). At the selected stages, half of the plants were transferred to +Fe, and the other half was exposed to - Fe. Fe treatments were started 3-4 hours after the growth chamber lights had switched on. -Fe hydroponic solution was the same quarter-strength Hoagland medium described above, with no Fe supplementation (0 µM FeNaEDTA). After 3 days of +Fe and –Fe treatment, roots, old leaves, young leaves and shoot apices were excised. Leaves were pooled according to the scheme shown in Figure S1C for the different growth stages. For leaf-by-leaf analysis, 3-week-old hydroponically grown plants were transferred at the RS to +Fe or -Fe condition, and each day leaves were harvested from day 1 to day 7 for leaf area analysis.

**Figure 1.**
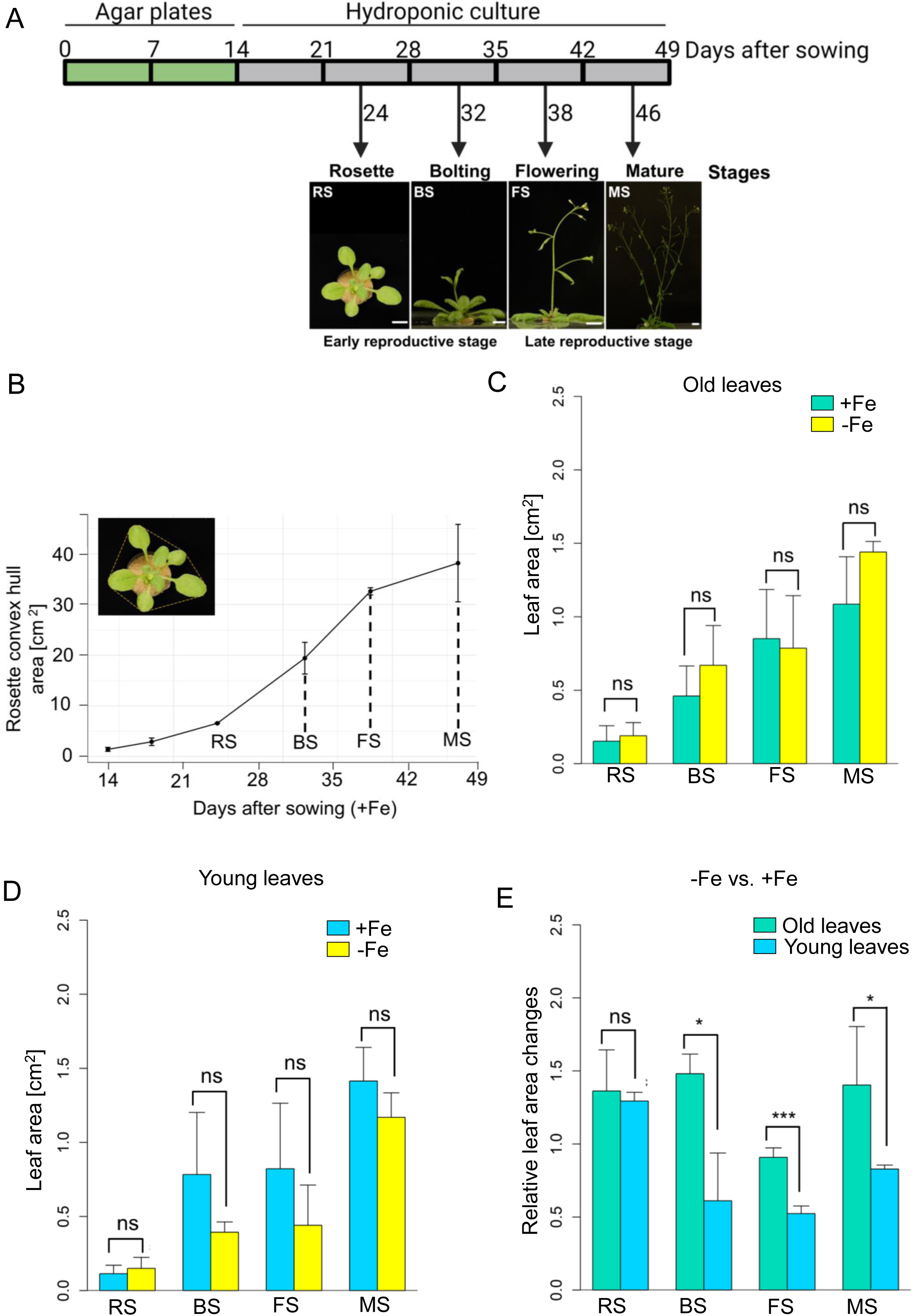
Growth dynamics during progression of the *Arabidopsis* life cycle. (A) Overview of experimental setup and reproductive growth stages selected for analysis of *Arabidopsis thaliana*. Seedlings were germinated on Hoagland agar plates before transplanting to standard hydroponic media with 25 µM Fe (+Fe) for growth to the four selected stages under long-day conditions; rosette stage (RS), bolting stage (BS), flowering stage (FS) and mature silique stage (MS) before further analysis, scale bar = 1cm. Further details on the reproductive growth stages in Figure S1A, B. (B) Rosette convex hull area increase of plants continuously grown on media with +Fe until the mature stage. The area included all leaves forming around the canopy plus their petioles (*n* = 2-8). (C-D) Absolute leaf areas of old and young leaves at +Fe and –Fe, respectively. Plants were transferred to +Fe and –Fe conditions for 3 days at the indicated growth stages. Old and young leaves harvested for analysis are described in Figure S1C. (E) Relative changes in leaf area of old and young leaves under –Fe compared to +Fe conditions at the stages RS to MS. Data in (C-D) represent average values of biological replicates (*n*=5). Error bars show ± SD. Statistical analysis was done by Student’s t-test (p < 0.05).

### Determination of rosette and leaf size

Plant rosette and leaf images were taken from the top using a Sony Alpha 6000 digital camera. Images were imported into ImageJ version 1.8.0 (https://imagej.nih.gov/ij/) for further processing and analysis. In brief, rosette images were converted to black and white and a color threshold set before quantification of rosette convex hull area. This was calculated from the surface area covered by the circumference of the longest leaves as shown in Figure 1B. Individual leaves were dissected from each plant and arranged in a row from oldest to youngest leaf as illustrated in Figure S1C. Leaf areas were determined by the area corresponding to that of the leaf blades.

### Scanning electron microscopy

Hand-dissected shoot apices were fixed overnight in 1 % paraformaldehyde (PFA) solution then dehydrated in a series of 30 %, 50 %, 70 % and 90 % ethanol, each for 20 min, and 100 % ethanol for 40 min. Samples were completely dehydrated to a critical point in a dryer (EMCPD 300, Leica) at 70 bar with 13°C /40°C cool-heat cycles for 3 hours. Samples were gold-coated under a high vacuum sputter (CCU-010, Safematic) before imaging with an electron microscope (Supra 40VP, Zeiss). Images were analyzed in ImageJ v 1.8.0.

### Gene expression analysis by reverse transcription (RT)-quantitative PCR (qPCR)

All steps of RT-qPCR analysis were conducted according to (Ngigi & Bauer, 2023). Briefly, plant materials were harvested and immediately deep-frozen in liquid nitrogen. Hand-dissected inflorescence shoot apices, however, were first placed in RNAlater (Thermofisher Scientific) and then deep-frozen. Samples were pulverized in a Precellys 24 tissue homogenizer (Peqlab, Germany). Total RNA extraction from all plant samples was performed using a plant RNA kit (Qiagen, Germany). Approximately 1 µg of RNA was treated with DNase I (Thermofisher Scientific) for removal of genomic DNA contamination followed by first-strand cDNA synthesis using the Revert-aid reverse transcriptase kit (Thermofisher Scientific). Real time qPCR was performed using 2x iTaq SYBR Green Supermix detection (Bio-Rad) on C1000 Touch CFX384 (Bio-Rad). qPCR data were individually checked and validated by melt curve analysis. Mean absolute normalized transcript abundances were calculated thanks to mass standard analysis and referral to the expression of reference gene *EF1B-ALPHA2*. qPCR primer sequences are listed in (Supplementary Table S1).

### Ionomics analysis

Roots and leaves were collected. Residual Fe in the root apoplast was removed by washing the roots in 20 mM Tris, 5 mM EDTA, pH 8 for 5 min before rinsing in deionized water. Old and young leaves were washed in deionized water to remove Fe on the leaf surface. All samples were dried at 65°C for 3 days (or longer as needed) before grinding to a fine powder. Ground material was transferred into metal-free 5 ml screw-cap tubes (Eppendorf, Germany, Cat. No. 0030122305) and 1 ml of 67 % nitric acid added. Samples were digested by boiling until a clear liquid was obtained. Elemental analysis was performed using inductively coupled plasma-mass spectroscopy (ICP-MS, Agilent 7700). A total of 5 plants were pooled together for material collection per sample in three independent experiments. Elemental concentrations were determined with the help of mass standards and normalized to sample dry weights.

### Statistical analysis and data visualization

Data was analyzed by one-way analysis of variance (ANOVA) with Tukey’s Honest Significant Difference (HSD) test for multiple comparisons and Student’s T test for pairwise comparisons. Bar graphs and line charts show mean and standard deviation of 3-5 biological replicates. Means that were statistically significantly different are indicated by different letters (p < 0.05). Correlation analysis of normalized gene expression values and mineral content was done using Pearson’s correlation with significance p value < 0.05. Dynamic correlation networks were visualized with the assistance of Julius AI (https://julius.ai/). Heatmaps were generated using mean absolute normalized gene expression values. The data were processed using R (version 4.4.1). Gene names were assigned as row labels. Z-score normalization was applied to each gene (row-wise) to account for variability in baseline expression. The Z-scores were calculated using the scale () function in R, with the following formula:

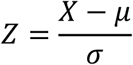

where X is the expression for a specific sample, μ is the mean expression of the gene across all samples, σ is the standard deviation of the gene expression. This row-wise Z-score normalization made it possible to compare expression levels of each gene across tissues at both +Fe and -Fe, allowing positive Z-scores to indicate above-average expression for high expression values and negative Z-scores to indicate below-average expression for low expression values. Unbiased gene clustering analysis was done by hierarchical clustering. The heatmaps were generated using the pheatmap package in R.

## Results

### The size of the leaf rosette increases during four reproductive phases

To assess plant responses during the reproductive phase, Arabidopsis wild-type (Col-0) plants were grown for up to seven weeks in sufficient Fe condition (+Fe) in a hydroponic system. Four meaningful reproductive stages were selected for plant phenotyping (Figure 1A, Figure S1A, B). At the rosette stage (RS, 24 days) plants did not yet have an emerged inflorescence stem, the rosette had approximately 11 emerged leaves, and plants had a closed inflorescence bud at the shoot apex (Figure 1A, Figure S1A-D). At the bolting stage (BS, 32 days), the main inflorescence stem had emerged and was between 2 to 4 cm tall, the rosette had around 11 examinable leaves which had grown bigger than at the RS, with closed floral buds (BS, Figure 1A, Figure S1A-D). At the flowering stage (FS, 38 days), the main inflorescence stem had elongated and was taller than 5 cm with 3-5 open flowers, the rosette had 13-17 amenable leaves (FS, Figure 1A, Figure S1A-D). At the mature silique stage (MS, 46 days), more than three lateral inflorescence branches had emerged, the main inflorescence stem was longer than 8 cm with 6-7 drying siliques, each being longer than 1 cm, and the rosette had more than 20 amenable leaves (MS, Figure 1A, Figure S1A-D). Rosette convex hull area is a commonly used parameter for growth evaluation of *A. thaliana* (Saini et al., 2022). The rosette growth curve was rather steep (Figure 1B). Maximal rosette dimensions were reached at MS. The fastest rosette size increase took place between RS and FS, particularly around BS, while it slowed down before the RS and between FS and MS (Figure 1B). We suspected that Fe was needed for rosette leaf growth to occur during the reproductive stages, particularly during the rapid size increase.

### Leaf areas of young leaves were reduced in response to -Fe versus +Fe

We tested whether leaf growth was linked with Fe availability. As according to a previous study, we termed the fully grown leaves that had reached their final or nearly final size as “old leaves”, while the still developing leaves were termed “young leaves” (Schuler et al., 2012). It is reported that the sizes of young leaves change upon micronutrient deficiency (Swartz et al., 2023). But whether rosette leaf growth is still influenced at the reproductive growth stages is not yet known. We grew plants up to the four selected reproductive stages and exposed them to +Fe and -Fe for 3 days. Individual leaves were dissected (Figure S1C) and leaf areas measured, then grouped into old and young leaves taking into account the leaf characteristics as described in (Swartz et al., 2023) (Figure S1C). There were no significant differences between either old leaves or young leaves at +Fe and -Fe, respectively (Figure 1C-D). However, when comparing the relative leaf area changes between -Fe versus +Fe, there were differences between old and young leaves at BS, FS and MS (Figure 1E), which can be attributed to an age-dependent expansion during early development phase. At the RS, old and young leaves had similar sizes when plants were exposed to +Fe and -Fe treatments (Figure 1D), presumably because they were still small and not expanding as much as during BS to MS.

To assess whether overall plant growth was also affected by -Fe treatment, we collected biomass data. The root fresh weights had increased steadily from RS to MS. At the RS and BS, there were no significant differences in root fresh weights between +Fe and -Fe treatment, however, at FS and MS, there was a drastic reduction of root fresh weight after the 3-day -Fe exposure compared with +Fe (Figure S1E). This finding potentially represents a survival strategy whereby root growth at late reproductive stages may slow down under Fe-limiting conditions due to Fe release in cells causing Fe toxicity (Naranjo-Arcos et al., 2017). Shoot biomass increased steadily from RS to MS. Particularly between FS and MS, shoot fresh weights had drastically increased, but there was no difference following the 3-day +Fe and -Fe conditions (Figure S1F). The shoot biomass increase was caused by the growing inflorescence stems, flowers and siliques, and perhaps the growth differences could not be detected by total weight.

Taken together, our growth system is suited to detect Fe-dependent growth dynamics in the rosette during the reproductive growth phases. In particular, the rapid increase of the sizes of young leaves and the rosette visible at the BS is influenced by Fe supply and suppressed in response to -Fe. Since the responses of rosette leaves during the reproductive phase are underexplored, we focused on rosette leaf physiology in the continuation of this study.

### Leaf-by-leaf analysis during consecutive days displayed a rapid growth arrest of young leaves in response to -Fe

To better resolve the Fe-dependent influence on leaf growth, we selected the RS stage and exposed plants to +Fe and -Fe. The rosette convex hull area and leaf areas of the first up to ten amenable leaves were determined on days 1, 2, 3, 5 and 7 (Figure 2). Between days 5 and 7, the rosette sizes of +Fe plants began to increase rapidly, and at these timepoints, significant differences became apparent between +Fe and -Fe-grown plants (Figure 2A-C). In the +Fe condition, rosette leaves displayed a green color throughout the 7-day period, while under -Fe conditions, the visual symptoms of leaf chlorosis became evident on day 2 and intensified as the experiment progressed (Figure 2A & B). Notably, -Fe leaf chlorosis was more pronounced in the young leaves, e.g. L8 to L10, compared to old leaves, e.g. L3 to L5 (Figure 2B). The young leaves e.g. leaf 8 were smaller at -Fe than +Fe from day 1 to day 7 (Figure 2D), similar to leaf 5 to leaf 7 (Figure S2D-F). L3, an old leaf, did not show changes in the leaf areas on day 1 and day 5 to 7 irrespective of treatments, showing that they had reached their final size at the RS. However, L5 to L8 increased in size from day 1 until day 7, indicating they were continuously developing young leaves (Figure 2D and S2D-F). Interestingly, already after 1 day of -Fe treatment, the leaf areas were smaller in all these four leaves (L5 to L8) compared to +Fe (Figure 2D and S2D-F). This shows that -Fe affected leaf growth and expansion immediately within a day.

**Figure 2.**
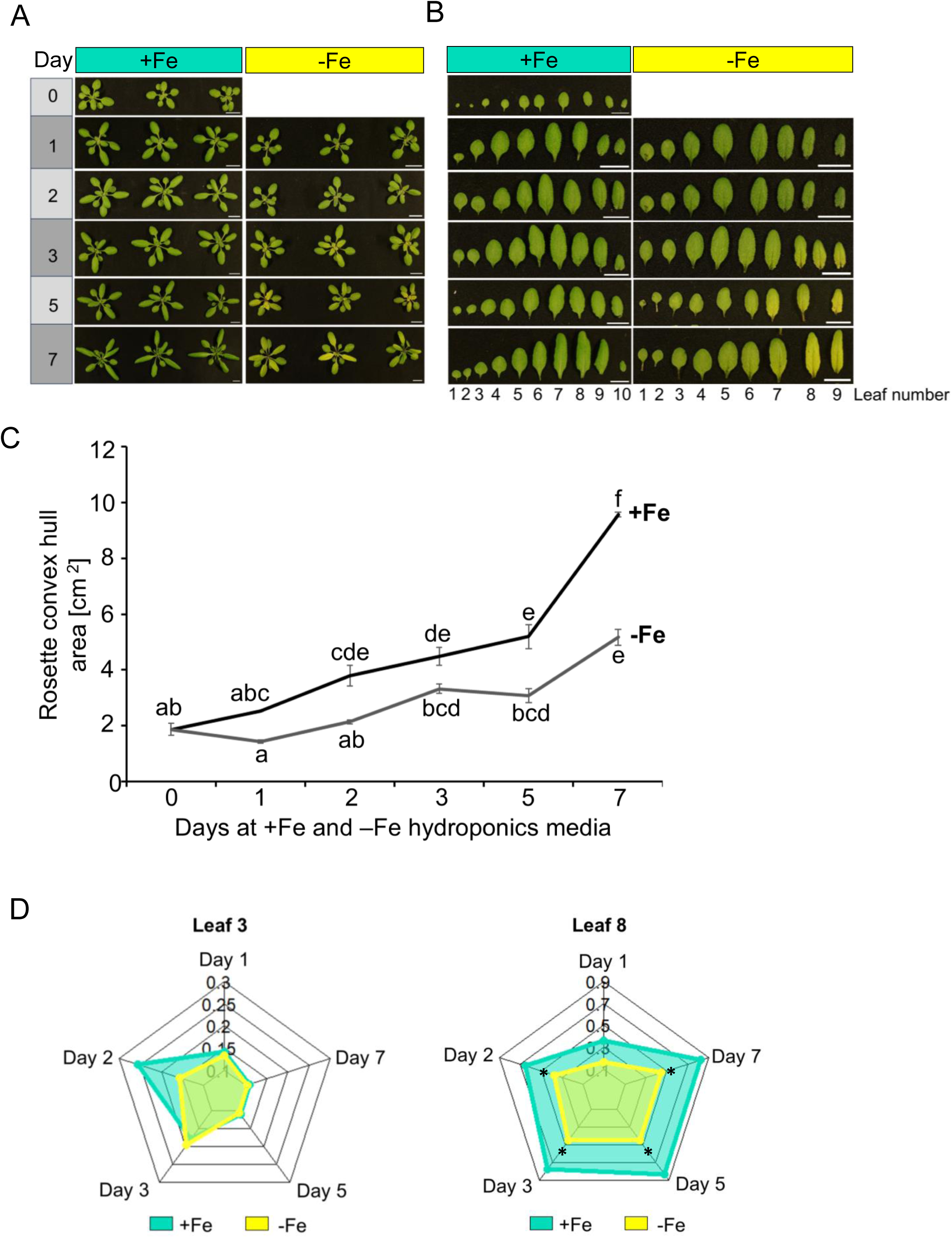
Leaf-by-leaf analysis of *Arabidopsis thaliana* at the rosette stage (RS). (A) At the RS, plants were transferred to respective Fe sufficient (+Fe) or deficient (-Fe) Hoagland media solutions and growth recorded from day 0 till day 7. (B) Leaf images of leaf 1 (L1) to 10 (L10) at the two Fe conditions, as indicated. (C) Cumulative rosette convex hull area as recorded from (A). Data represent means of biological replicates (*n*=3). Error bars represent ± SD. Statistical analysis was done using one-way ANOVA and Tukey’s HSD test. Means followed by the same letter are not significantly different (p < 0.05). (D) Radial charts showing the leaf areas of leaf 3 and 8 after exposure to either +Fe (control) or –Fe conditions from day 1 to day 7. Radial charts were plotted in R Studio using “fsmb” package. Plants were grown for 3 weeks before culturing in either +Fe or -Fe conditions. Asterisks show where leaf areas were statistically significantly different between +Fe and -Fe conditions, (p<0.05), one-way ANOVA and Tukey’s HSD test.

Taken together, these findings underscore the critical role of Fe for the growth of young leaves. Young leaves were more susceptible to developing leaf chlorosis than old leaves. The differential phenotypic responses observed between old and young leaves speak in favor of differential Fe homeostasis physiology between old and young leaves. The very rapid growth response indicates that -Fe sensing and signaling directly controls leaf growth.

### Expression of Fe-regulated genes points to Fe recycling taking place in old leaves at RS and BS

To capture differential regulation patterns of leaves across the reproductive stages, we studied the gene expression patterns of 23 Fe homeostasis genes by analyzing transcript amounts using reverse transcription-qPCR, that corresponded to meaningful lists of genes from previously reported studies (Mai et al., 2016; Schwarz & Bauer, 2020). We grew plants as above and collected old and young rosette leaves of +Fe and -Fe plants across the four reproductive stages to investigate gene expression in space- and time-resolved manner (old leaves were L3 to L5 from RS to FS, and L5 to L7 at MS, as L1 to L4 had senesced at this stage; young leaves were L9 to L11 from RS to FS, and L14 to L17 at MS; Figure S1C). Inflorescence shoot apices were collected since they harbored the youngest leaves and leaf primordia. Roots were harvested to estimate the Fe status of plants. The 23 analyzed genes were grouped into six categories or clusters according to their previously described co-regulation and expression patterns, representing different roles in Fe homeostasis (Figure 3A).

**Figure 3.**
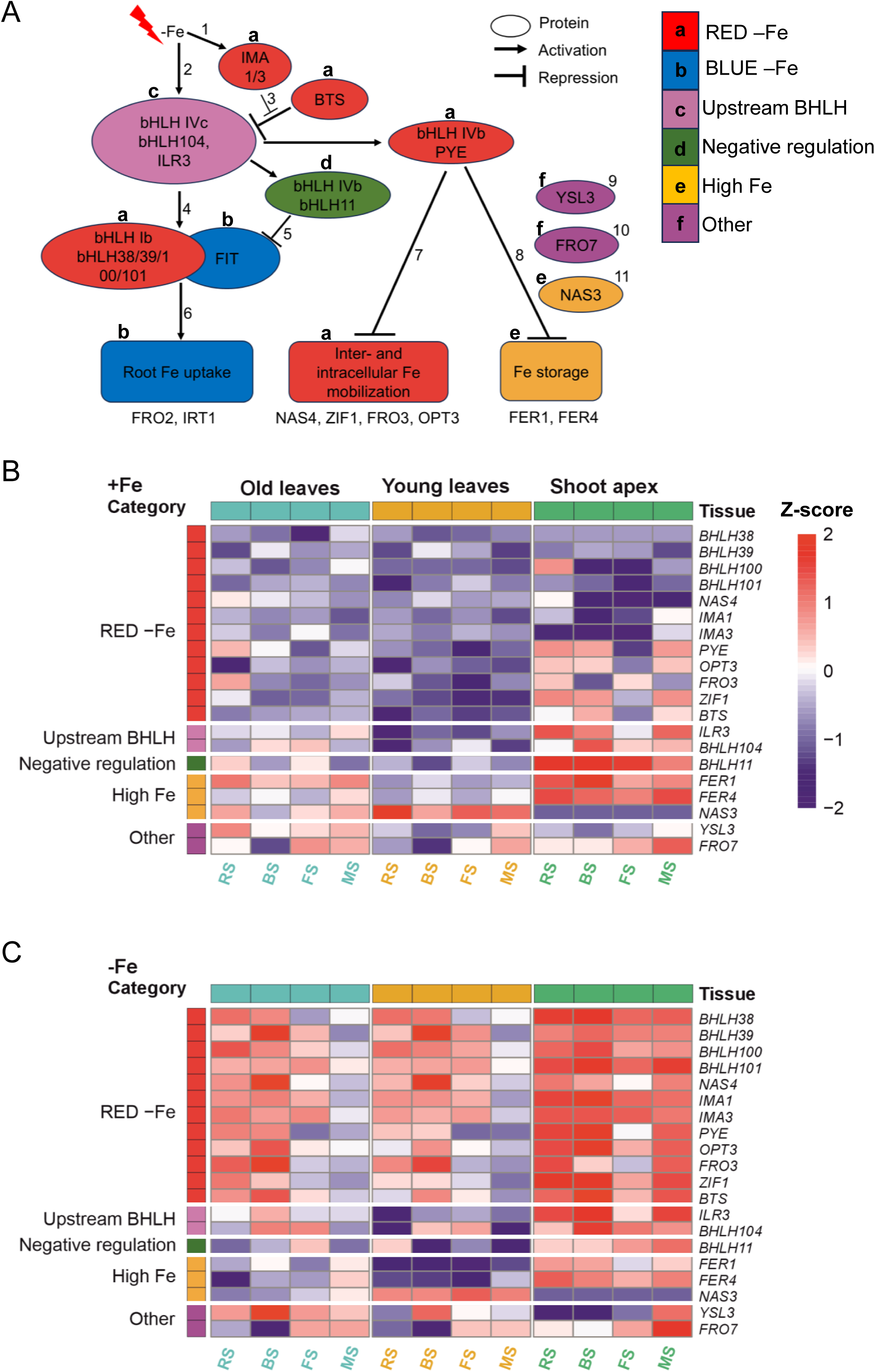
Gene expression analysis of selected Fe homeostasis genes. (A) Schematic illustration showing connections and functions of Fe homeostasis genes used in gene expression analysis. Represented is the Fe-regulated transcriptional cascade and multiple layers of co-expressed genes involved in plant Fe homeostasis. Exemplary references: ^1^(Kobayashi et al., 2021), ^2^(Zhang et al., 2015), ^3^(Lichtblau et al., 2022), ^4^(Li et al., 2016), ^5^(Tanabe et al., 2018), ^6^(Trofimov et al., 2019), ^7^(Long et al., 2010), ^8^(Tissot et al., 2019), ^9^(Chu et al., 2010), ^10^(Mukherjee et al., 2006), ^11^(Schuler et al., 2012). Letters above the genes correspond to the co-expressed gene categories. (B-C) Heatmaps showing the absolute expression profiles of 20 selected Fe homeostasis genes in old leaves, young leaves, and inflorescence shoot apex after exposure to +Fe and –Fe for 3 days. Plants were grown to the various stages, as indicated, before transfer to either Fe solutions. The growth stages RS, BS, FS and MS correspond to rosette stage, bolting stage, flowering stage and mature stage, respectively. Heatmaps were generated in R studio version 4.4.1 using calculated z-scores of normalized gene expression values of each gene across tissues for both +Fe and –Fe. Functional gene category annotations are displayed on the left side of the heatmap, with tissue-type annotations displayed on the top. Gene upregulation is represented in red colour and gene downregulation with blue colours, p < 0.05 (n = 3). The colour scale ranges from -2 to 2 to represent the variation in gene expression.

The gene expression data were remarkable in several ways. At first, we investigated gene expression in the leaf and shoot apex samples. To get an overview of how the genes are expressed across the four stages and tissues, we performed a principal component analysis (PCA) of all genes. Results revealed that the FIT-independent RED -Fe gene cluster separately grouped from other gene categories, highlighting differential regulation by Fe (Figure S3). The RED -Fe cluster genes which serve the recycling of Fe in the whole plant were overall expressed more under -Fe versus +Fe in both, the old and young leaves, and the shoot apex samples (Figure 3B, C). In the leaf samples, this was particularly evident at the early reproductive stages RS and BS where all 12 genes in the RED -Fe cluster were expressed at higher level than at late reproductive stage, MS (Figure 3C). We see this observation as a hint that the -Fe-induced RED cluster genes are not needed to be expressed strongly in the rosette leaves at the late reproductive stages when siliques are developing. We suspect that at FS and MS the rosette leaves become less susceptible to -Fe signals than at earlier stages. Surprisingly, gene expression of RED -Fe cluster was remained induced in the inflorescence shoot apices at FS and MS, which provided a hint that these genes regulated by Fe homeostasis in the reproductive shoot apical meristem. We noted another interesting feature about the expression patterns of RED -Fe cluster genes, namely that their co-expression was split in the +Fe samples. The four *BHLH* subgroup Ib genes (*BHLH038*, *BHLH039*, *BHLH100* and *BHLH101)*, *NAS4*, *IMA1,* and *IMA3* genes were expressed at comparably low levels at +Fe as compared with *PYE*, *OPT3*, *FRO3*, *ZIF1* and *BTS*, which were more expressed at +Fe and even quite elevated in the +Fe shoot apices. Under -Fe, hierarchical clustering revealed that RED-Fe genes were differentially upregulated and co-expressed by Fe deficiency, suggesting co-regulation by Fe in shoot samples (Figure S4). In sum, the -Fe regulation of RED -Fe cluster genes is most important in both old and young leaves, in particular, at the BS and RS, as well as in the inflorescence shoot apices at all reproductive stages. There is evidence that *PYE*, *OPT3*, *FRO3*, *ZIF1* and *BTS* are differently expressed from other RED -Fe cluster genes, which can hint to different upstream regulation or functions.

Next interesting observation was that the expression of genes encoding the positive regulators of RED -Fe cluster genes, namely *ILR3* and *BHLH104*, summarized as Upstream BHLH, and the negative regulator, namely *BHLH11*, was consistent with that of the RED -Fe cluster genes *PYE*, *OPT3*, *FRO3*, *ZIF1* and *BTS*. When these RED -Fe cluster genes were up-regulated specifically at BS under -Fe, it was also the case for the positive regulators *ILR3* and *BHLH104*, while the negative regulator *BHLH11* was down-regulated, especially in the old leaves. The upstream regulator genes were not Fe-regulated and were expressed at BS to MS in old and young leaves, and in all the stages in inflorescence shoot apices (Figure 3B, C). For *ILR3* and *BHLH104*, this was expected, but for *BHLH11* it was unexpected, since it was reported that BHLH11 is down-regulated by -Fe (Tanabe et al., 2019). With respect to a possible difference of expression in old versus young leaves, we noted a tendency that at -Fe, expression was higher in old versus young leaves in agreement with the expression of target RED -Fe cluster genes (Figure 3C).

Another remarkable point was that High Fe marker *FER1*, a marker gene for Fe sufficiency and excess conditions (Briat et al., 2010; Roschzttardtz et al., 2013), expression levels remained relatively more highly expressed throughout the stages at +Fe in the old leaves compared to -Fe and young leaves (Figure 3B). This was not the case for *FER4* expression. On the other hand, *FER1* and *FER4* showed similar high expression in the inflorescence shoot apex samples across the stages at +Fe with slight downregulation at -Fe (Figure 3B, C). Surprisingly, *NAS3* expression levels was consistently up in the young leaves at all the four reproductive stages in both Fe conditions, but down in shoot apices (Figure 3B, C). With respect to the question about differences in the leaf samples at -Fe, we noted tendencies of higher expression of *FER1* and *FER4* in old versus young leaves, both of which were significant at MS whereas *NAS3* was higher expressed in young versus old leaves across the four stages (Figure 3C). Thus, High Fe marker gene expression did not distinguish the four reproductive stages, but it might be a marker for Fe-regulated processes in the old versus young leaves and in the shoot apex. Perhaps *NAS3* plays a major role in Fe remobilization and export.

Finally, *YSL3* was expressed at higher levels in leaves than in the shoot apex samples, and in some cases, there appeared up-regulation at -Fe versus +Fe, such as at the BS in leaves (Figure 3B, C). *FRO7*, had similar expression between +Fe and -Fe, but it was mostly expressed at higher levels in shoot apex samples than in leaves, especially at MS (Figure 3B, C). Very interestingly, the expression of *YSL3* was up-regulated in old versus young leaves at -Fe at all the four stages RS, BS, FS and MS. *FRO7* expression differences was not noted between leaves and Fe supply (Figure 3B, C).

The root gene expression patterns were also very remarkable. As compared with rosette leaves and shoot apex samples, the expression levels were surprisingly lower for most genes analyzed above (Figure S5). Although the FIT-dependent BLUE -Fe genes, previously known as root -Fe markers, were on one side expressed at similar or higher levels in roots versus the shoot organs, but on the other side, an exception was that the BLUE -Fe genes were quite strongly expressed at BS in the leaf samples, and at MS in the shoot apex samples. Perhaps the root system is either no longer very sensitive to -Fe or it also consists of different zones that respond differently to -Fe.

Taken together, the leaf and shoot apex gene expression patterns were rather complex during the reproductive stages. There were differences in expression between +Fe and -Fe, between the four reproductive stages and between old and young leaves as well as inflorescence shoot apex. Two major conclusions could be drawn. Fe recycling genes of the RED -Fe cluster were mostly expressed at the RS and BS stages in old and young leaves at -Fe, while High Fe markers were rather uniformly expressed throughout all stages at both +Fe and -Fe. Most genes that discriminated against old and young leaves were expressed at higher levels in old versus young leaves, especially RED -Fe cluster genes *PYE*, *OPT3*, *FRO3*, *ZIF1* and *BTS* and *YSL3* at -Fe, as well as *FER1* at +Fe and in the High Fe genes cluster. Only *NAS3* was found to be consistently higher expressed in young than old leaves and shoot apices. These findings indicate that different physiological states are associated with the organs at the different reproductive stages.

### Mineral contents differ between old and young leaves at the early reproductive stages

To test whether old and young leaves may be subject of different Fe and metal ion physiology, we determined the elemental contents of Fe, Zn, Mn and Cu.

At first, we compared the elemental contents between +Fe and -Fe separately for old and young leaves and roots (Figure 4). In old and young leaves, there was a clear trend that metal contents decreased from RS to MS in the +Fe and -Fe conditions (Figure 4A, B, D, E, G, H, J, K). In old leaves, this was the case for Zn and Mn at +Fe and -Fe (Figure 4D, G) and for Cu at -Fe (Figure 4J). In young leaves, it was the case for all elements, however, in a Fe-dependent manner, such as for Fe and Mn at +Fe (Figure 4D, G), and for Zn and Cu at both Fe supply conditions (Figure 4E, K). We then compared the elemental contents at each stage between +Fe and -Fe in leaves. Surprisingly, there were not many differences in the old leaves when we compared across the stages +Fe and -Fe conditions. In old leaves, there were no differences in Fe, Zn and Mn contents between +Fe and -Fe treatments and across the growth stages (Figure 4A, D, G). However, there was a difference in Cu contents, which had increased at RS in the old leaves (Figure 4J). In young leaves, on the other hand, the Fe contents had clearly decreased between +Fe and -Fe from RS to FS (Figure 4B). The Cu contents had again increased between +Fe and -Fe at the RS (Figure 4K). When comparing old and young leaves, the metal contents were either unchanged or lower in young versus old leaves in the cases of Fe, Zn and Mn (Figure S6A-F). Only Cu amounts were higher in young than in old leaves at BS under +Fe condition (Figure S6G). Hence, unlike young leaves, it seems that Fe was likely not mobilized from old leaves during the reproductive stages at -Fe, and Fe was also not likely moved from old to young leaves. This was different for Mn and Zn, which were likely mobilized in old and young leaves as their contents decreased with progression of reproduction.

**Figure 4.**
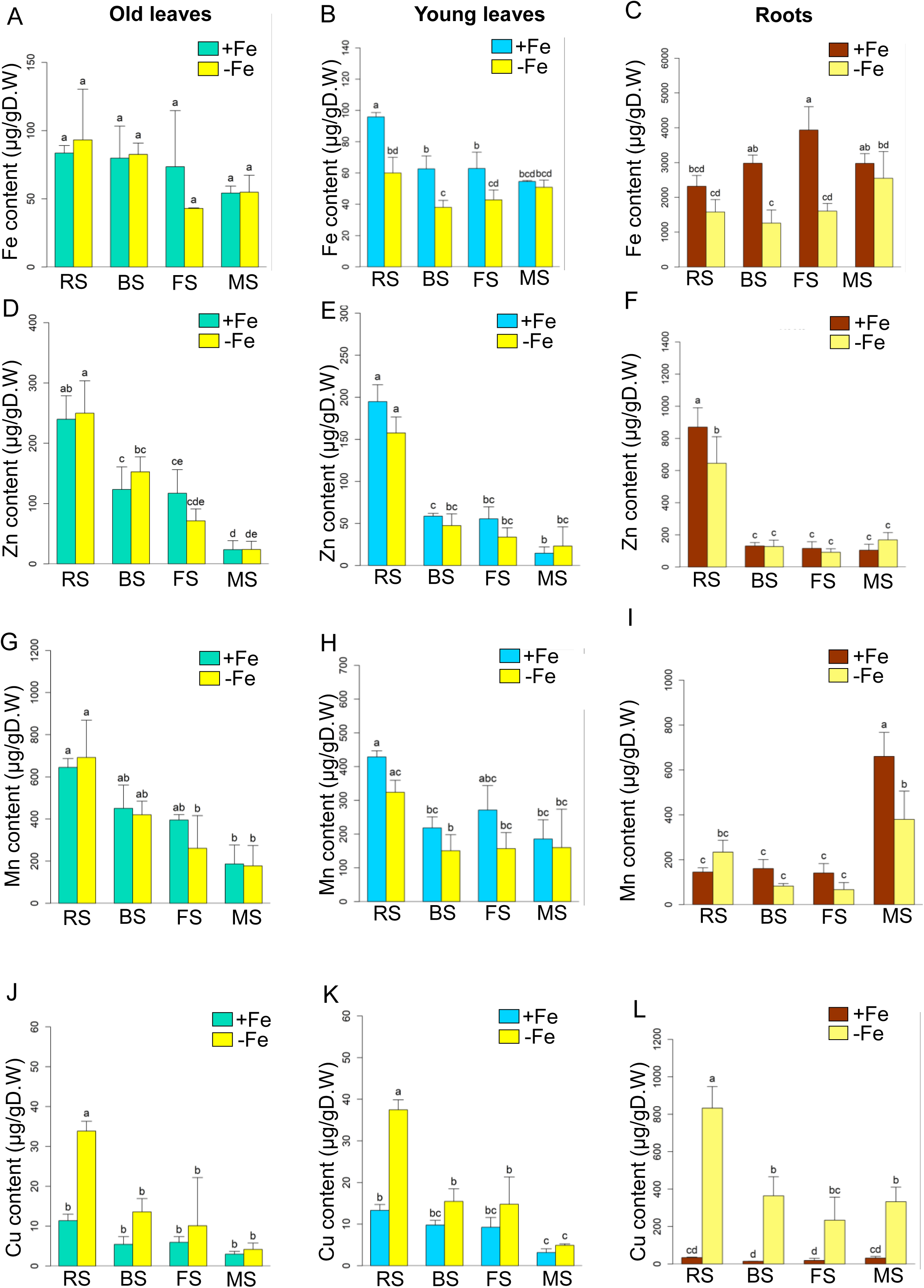
Ionomic analysis of leaves and roots. Fe (A-C), Zn (D-F), Mn (G-I), and Cu (J-L) content per dry weight (D.W.) in old leaves, young leaves and roots of plants grown at 25 µM Fe (+Fe) or 0 µM Fe (–Fe) conditions and at the four growth stages; RS, BS, FS and MS. Plants were exposed to 3 days of +Fe or –Fe before sample collection. Data represents mean of three biological replicates (*n* = 3). Error bars represent ± SD. Letters on top of each bar show level of significance according to one-way ANOVA and Tukey’s HSD test, p < 0.05.

The root elemental contents supported that root uptake was dependent on Fe supply. As expected, Fe contents were lower in roots at -Fe versus +Fe, even though this only occurred at BS and FS (Figure 4C). Unexpectedly, the content of Zn was lower at RS and that of Mn was lower at MS when comparing -Fe against +Fe (Figure 4F, I). Notably, the Cu content was up in -Fe versus +Fe roots at all stages (Figure 4L), which presumably accounts for the elevated Cu contents in -Fe leaves at RS.

Hence, complex interactions in metal ion physiology were observed in old and young leaves at +Fe and -Fe across the stages. -Fe affected Fe, Zn and Mn contents similarly, but had very different effects on Cu contents both in leaves and roots. The highest metal ion contents were mostly observed at the early reproductive stages, indicating a usage during progression to late reproductive development.

### Gene expression correlated with the mineral contents in leaves

We investigated whether the gene expression levels correlated with mineral contents. Notably, we observed positive correlations between Fe, Zn, Mn and Cu levels, in both old and young leaves at +Fe and -Fe conditions. This confirmed an interaction in the mineral metal physiology in both leaf types under changing Fe environments. Furthermore, we found meaningful correlations between mineral metal contents with gene expression levels in old and young leaves (Figure 5, S7).

**Figure 5.**
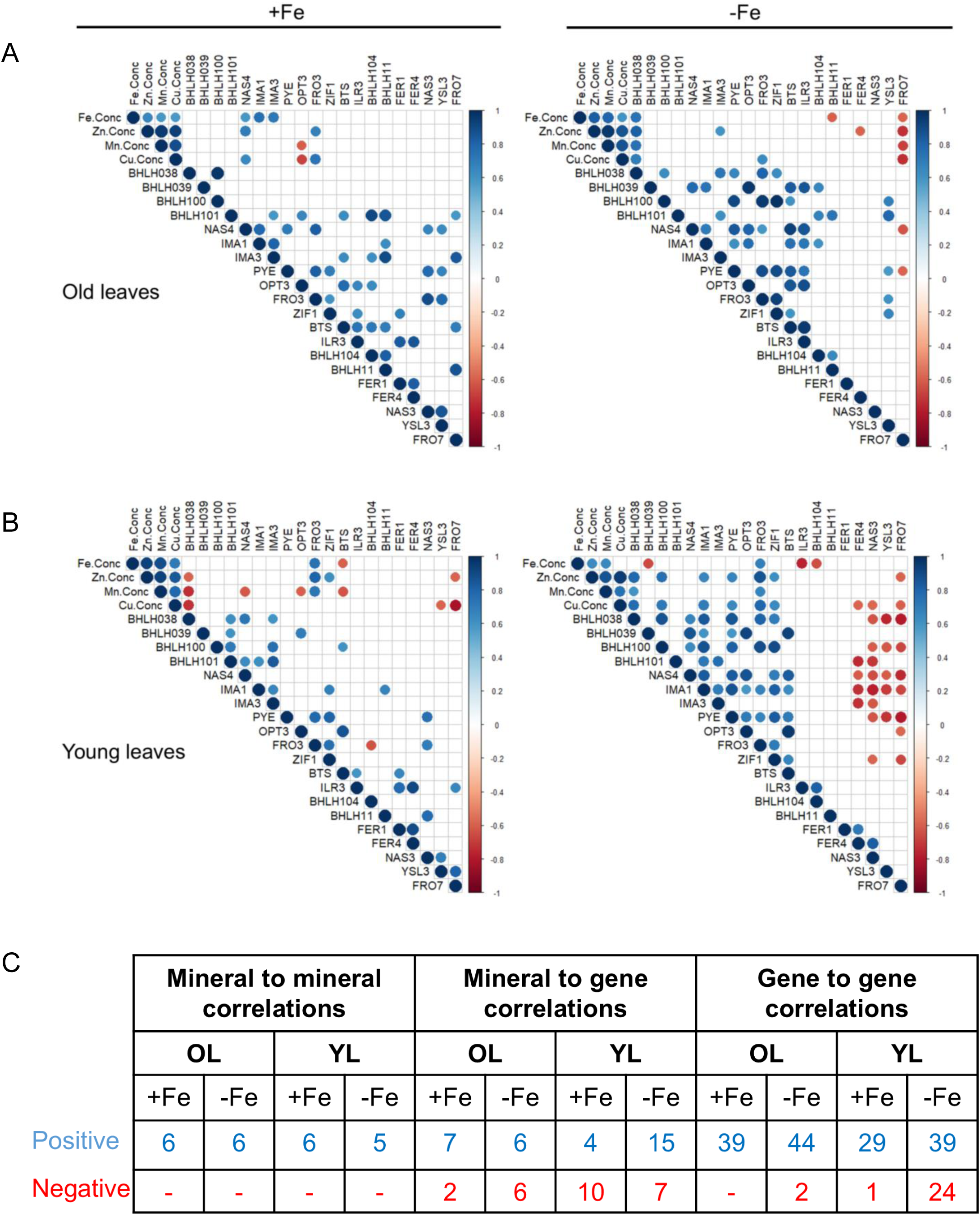
Correlation analysis between gene expression changes and ionomic contents. Correlations were performed with data from old leaves (A), and young leaves (B) at +Fe (right side) and -Fe (left side) conditions. Analysis was done using averages of normalized gene expression values and mineral concentrations of Fe (Fe Conc.), Zn (Zn Conc.), Mn (Mn Conc.) and Cu (Cu Conc.) across all four stages RS, BS, FS, and MS. Color scale represents Pearson’s correlation coefficients normalized in the range between -1 to 1, whereby -1 is negative correlations and 1 is positive correlations calculated between genes and mineral content pair. Significant correlations at p<0.05 are displayed in the diagram. (C) Summary table showing number of correlations between either mineral-to-mineral content, mineral content to gene expression level and correlation of gene gene-to-gene expression level in old and young leaves. Numbers show total number of positive and negative correlations at both +Fe and –Fe conditions.

In old leaves, Fe showed most significant correlations with genes at +Fe while Zn was the most correlated in comparison to other minerals at -Fe (Figure S7A-B). Fe contents correlated positively with the three coexpressed RED -Fe genes *NAS4*, *IMA1* and *IMA3* at +Fe, and with *BHLH038* at -Fe (Figure 5A, S7A-B). Negative correlations were found at -Fe between Fe content and *BHLH11* and *FRO7* expression. Furthermore, in old leaves, Zn and Cu content correlated positively with expression of RED -Fe genes *NAS4* and *FRO3* at +Fe, and *BHLH038* at -Fe (Figure S7A-B). On the other hand, *OPT3* negatively correlated with Mn and Cu levels under +Fe condition. Positive correlations were seen between Zn, Mn and Cu amounts with *BHLH038* at -Fe, and negative correlations of these minerals content with *FRO7* expression (Figure 5A and S7B).

In young leaves (Figure 5B), different correlations were seen, and most correlations were seen in young leaves at -Fe. Interestingly Mn, Zn and Cu were most correlated with genes at +Fe whereby Cu showed most correlations under -Fe conditions (Figure S7C-D). Fe contents correlated positively with *FRO3*, and negatively with *BTS* at +Fe (Figure 5B, S7C). At -Fe, the Fe amounts positively correlated with *FRO*3, and negatively with *BHLH039, ILR3 and BHLH104* gene expression (Figure 5B, S6D). There were positive correlations between Zn and Mn content and *FRO3* at +Fe . At similar Fe condition, negative correlations appeared between Zn and Cu with *BHLH038* and *FRO7*, while Mn negatively correlated with *BHLH038, NAS4, OPT3* and *BTS* expression levels. When Fe was absent, more positive correlations were observed between Zn content with *BHLH038/100, IMA1, PYE, FRO3 and ZIF1*, but negatively with *FRO7* (Figure 5B, S7D). Similar correlation patterns were observed with Cu amounts in the young leaves at -Fe. Additionally, Cu amounts showed negative correlations with *FER4, NAS3 and FRO7*. Similar to the old leaves, the contents of Fe, Zn, Mn and Cu in young leaves, positively correlated with each other at both Fe conditions, except between Fe and Cu at -Fe. A quantification of correlation between gene expression and mineral contents is shown in Figure 5C.

In summary, the observed correlation patterns confirmed that the gene expression was indeed linked with the mineral metal contents in the old and young leaves under +Fe and -Fe conditions. It was also possible to distinguish old and young leaf Fe physiology by the number of correlations among gene expression and metal contents. The mineral metal levels correlated with each other and may account for the observed gene expression changes in the leaves.

## Discussion

In this study, we investigated how reproductive transitions impact Fe and metal ion physiology in rosette leaves when plants were exposed to +Fe and -Fe. Leaf growth especially between RS and FS was markedly influenced by Fe availability, which indicates an interesting cross-connection with leaf signaling events influencing leaf growth. Contrary to our idea that -Fe would cause important mobilization of Fe from rosette leaves during late reproductive stages, our findings revealed unexpected stage-specific dynamics in Fe and metal ion homeostasis. At the early reproductive stages, plants demonstrated a stronger response to -Fe in the rosette leaves than at late reproductive stages, suggesting that rosette leaves at late reproductive stages do not play important roles in metal ion homeostasis. Old and young leaves reacted differently in terms of changes of metal contents and gene expression patterns in response to +Fe and -Fe. The data indicate that old and young leaves can be sources and sinks for metal ions under different Fe conditions.

### Growth of young leaves is rapidly inhibited at -Fe suggesting an intersection of -Fe and leaf growth signaling

Young leaf expansion was notably slowed and accompanied by leaf chlorosis following a 3-day - Fe treatment, particularly during the rapid growth at the BS and FS. The inhibitory effect of -Fe on growth and greening of young leaves was very rapid and became visible already after one day of -Fe. This fast Fe-dependent leaf growth response demonstrates the high plasticity of young leaves in response to environmental cues, in this case -Fe signals. Old leaves, instead, had already reached their final size so that their growth response was unaffected. Leaf growth is the result of cell proliferation and cell expansion (Li et al., 2024; Wu et al., 2009; Xu et al., 2016). We therefore suppose that the observed reduced leaf areas of young leaves could be a consequence of repression in growth and plasticity in these leaves upon -Fe signals. Future studies will focus on dissecting the players at the intersection between regulation of leaf growth and Fe homeostasis control.

It is known that leaf form and morphology change in response to environmental and developmental cues and are especially common for the juvenile to adult leaf transition, which is caused by a process referred to as leaf heteroblastic reprogramming (Bäurle & Dean, 2006; Costa et al., 2012; Tsukaya, 2005; Tsukaya et al., 2000). In *Arabidopsis thaliana,* SQUAMOSA PROMOTER BINDING PROTEIN-LIKE 9 (SPL9) targeted by miR156 regulates the juvenile-to-adult transition which influences leaf morphology (Wu et al., 2009; Xu et al., 2016). Mechanisms for leaf development transition and Fe homeostasis have been proposed for the miR156-SPL9 module, whereby SPL9 transcriptionally could regulate Fe deficiency markers *FIT* and *PYE* by direct promoter binding (Wang 2015), a model requiring further investigation. Furthermore, in rice, *OsSPL9*, an ortholog of *SPL9* in *Arabidopsis*, has been proposed to be involved in Cu deficiency response whereby a knockout *spl9* mutant was sensitive to Cu deficiency with reduced accumulation of Cu in the old versus young leaves (Wang et al., 2024). These studies confirm that indeed SPL9, among other leaf growth regulators, may influence mineral uptake responses to synchronize development with nutrient availability. Thus, it is possible that in our system, -Fe signals influenced expression levels of leaf growth regulators. Future studies may investigate Fe responses using mutants defective in leaf cell proliferation and elongation, for example, using plants with *SPL9* mutations which were shown to have defects in leaf cell expansion during the juvenile-to-adult transitioning (Li et al., 2024).

On the other hand, retrograde chloroplast signaling can also affect cell expansion and leaf greening (Andriankaja et al., 2012). Inferring from this, it is also a possibility that in our system, -Fe signals acted to inhibit leaf cell expansion and greening by blocking retrograde signaling. In such a scenario, -Fe signaling to the chloroplasts repress signals towards the nucleus and inhibit expression of nuclear-encoded leaf development- and photosynthesis-related genes. Previous studies have addressed retrograde signaling upon -Fe (Balparda et al., 2020, 2021). Alternatively, retrograde chloroplast signals may be activated under -Fe but then block the reprogramming in the nucleus thus influencing downstream transcription of photosynthesis and growth regulators (Nam et al., 2021). -Fe-induced transcription factors including the class 1b bHLH might be able to interact with the transcriptional regulation apparatus for leaf cell differentiation, thus regulating leaf cell proliferation and expansion as demonstrated before (Andriankaja et al., 2014). TCP transcription factors, that positively control leaf expansion can bind with - Fe-induced bHLH subgroup Ib proteins and transcriptionally repress their expression (Andriankaja et al., 2012). However, in these studies, bHLH subgroup Ib transcription factors seemed to be induced during the transition phase from cell proliferation to cell expansion at +Fe in early leaf growth phases. This appears contradictory to our -Fe data where these genes were induced in old leaves as well. Nevertheless, it needs to be taken into account that these previous studies investigated seedling stages while we focused on later reproductive stages which may account for such differences.

### Distinct Fe physiology in early and late reproductive stages and in old and young leaves

Rosette leaves display more distinctive Fe physiology responses at +Fe at the RS and BS than at late reproductive stages. The gene expression patterns and metal contents indicate that at the early reproductive stages, metal ion physiology is more active than at late reproductive stages. For example, Fe recycling genes (RED -Fe genes) *NAS4*, *OPT3*, and *FRO3* which play a key role in Fe redistribution between source and sinks (Khan et al., 2018; Kumar et al., 2017; Nguyen et al., 2022; Schuler et al., 2012; Stacey et al., 2008), were upregulated in both old and young leaves at RS and BS. This showed that Fe mobilization responses in old and young leaves were highly induced during these stages. Studies have demonstrated that Fe fluxes increase between the rosette and the inflorescence organs including stems, fruits and seeds after onset of flowering (Mendoza-Cózatl et al., 2014; Pottier et al., 2019; Waters & Grusak, 2008; Zhai et al., 2014). On the other hand, High Fe genes, which are upregulated by Fe sufficiency and excess conditions and downregulated by Fe deficiency (Briat et al., 2010; Roschzttardtz et al., 2013) like *FER1* and *NAS3* were expressed similarly throughout the four stages in old and young leaves respectively, while a marker for internal Fe mobilization from the chloroplast, *FRO7*, was predominantly expressed at the FS and MS stages in both leaves. This can indicate that Fe recycling from subcellular storage involving the RED -Fe genes is more important at the early reproductive stages. At these stages, old leaves can be sources of Fe and other metal ions for supply to sink organs including developing leaves and inflorescence stems. At the later reproductive stages, Fe remobilization from plastids involving genes like *FRO7* may be switched on. Fe is mobilized from plastids at the onset of leaf senescence and may prepare for Fe allocation towards seed filling (Himelblau & Amasino, 2001). Senescence takes place in the older leaves and can be triggered by plant age and environmental cues in order to enhance plant resilience to stress conditions and efficiently use nutrients (Buchanan-Wollaston, 1997; Lemaître et al., 2008; Nooden et al., 1996; Sperotto et al., 2012; Waters et al., 2009). Leaf senescence can lower the sensitivity to external -Fe as plants instead mobilize intracellular Fe stores in the vegetative organs instead of *de novo* uptake by the roots (Pottier et al., 2019). Additionally, resulting oxidative stress responses due to internal Fe release may induce expression of High Fe gene cluster in leaves consequently dampening Fe uptake responses in roots (Le et al., 2016). Rosette leaves are the targets of xylem mineral transport. There was a drastic decrease of Mn and Zn in old and young leaves and a decrease of Fe and Cu contents in young leaves at +Fe from RS towards MS, indicating that at late reproductive stages metals may have been exported from the rosette leaves to replenish sinks forming in the inflorescence such as fruits and seeds. It is likely that at the late reproductive stages the xylem stream is oriented towards the inflorescence stem directly rather than the rosette leaves so that cauline leaves and inflorescence stems account primarily for allocation of Fe, taken up de novo and transported from roots to seeds (Pottier et al., 2019; Schuler et al., 2012; Waters & Grusak, 2008). The enhanced -Fe responsiveness in the shoot apex samples even at FS and MS speaks in favor of this. In future studies it can be investigated whether new Fe storage tissues arise during reproductive growth in addition to the rosette leaves, e.g. in the secondary phloem tissues coupled with secondary growth in the Arabidopsis hypocotyl during flowering (Lehmann & Hardtke, 2016; Wunderling et al., 2018).

Notably, particular observations were highly interesting and striking. One remarkable observation was that the differential expression of RED -Fe genes in the old leaves at -Fe indicated a subfunctionalization among RED -Fe gene functions. The expression regulation of subgroup Ib *BHLH*, *IMA* and *NAS4* genes could be distinguished from that of *PYE*, *OPT3*, *FRO3*, *ZIF1* and *BTS*. PYE negatively regulates certain RED -Fe genes, including *bHLH1B* genes, *FRO3*, *ZIF1* and *NAS4* in seedlings (Haydon et al., 2012; Long et al., 2010; Pu & Liang, 2023; Schuler et al., 2012). Possibly, individual regulator proteins such as PYE, BTS or upstream transcription factors have individual functions in controlling positively or negatively different subsets of Fe mobilization and Fe storage genes in different tissues (Akmakjian et al., 2021; Muhammad et al., 2022; Rodríguez-Celma et al., 2019; Zhang et al., 2015) and at different developmental stages. Future studies can focus on investigating the roles of regulatory genes not only during early vegetative seedling stages but also during reproductive stages.

The expression of FIT-dependent genes in roots was induced in the old leaves, young leaves and shoot apex from the RS to MS with -Fe conditions. This shows age-dependent upregulation, especially with progression from early to late reproductive development. Indeed, reports have shown evidence of mRNA transport between shoots and roots, for example, *IRT1* was among the long distance transported transcripts in grafted plants grown under nutrient limitation (Thieme et al., 2015). BLUE -Fe genes can be expressed in shoots, even though this has not yet been explored. It has been shown that *IRT1* and *FRO2* are expressed in seedling leaves, even though to a less extent compared to roots (Jakoby et al., 2004). Moreover, *IRT1* promoter-driven reporter activity was observed in flowers (Vert et al., 2002). Furthermore, overexpression of FIT led to ectopic expression of *IRT1* and *FRO2* in leaves at +Fe and -Fe conditions (Meiser et al., 2011). This was also demonstrated in BHLH39Ox plants whereby *FIT* expression was evident in leaves of seedlings, and surprisingly to a similar level as wildtype plants (Naranjo-Arcos et al., 2017). This indicates that FIT and its direct target genes, *IRT1* and *FRO2*, are expressed in shoots, too. Thus, there is a possibility that BLUE -Fe genes are induced in leaves and reproductive meristems during later reproductive stages. Future studies should further explore the role of this age-dependent expression pattern. Another observation was interesting, namely the increase of Cu contents in old and young leaves at the RS following the -Fe treatment. This Cu increase was certainly linked with the very high Cu uptake into roots at -Fe. It indicates that despite their growth inhibition, the young leaves accumulated high Cu amounts at this stage. There are evidences for Fe and Cu crosstalk whereby Cu amounts in roots and shoots are increased at -Fe, and vice versa (Bernal et al., 2012; Carrió-Seguí et al., 2019; Chia et al., 2023; Kastoori Ramamurthy et al., 2018; Rai et al., 2021; Waters & Armbrust, 2013). One of the genes encoding a transporter for Cu uptake is *COPT2* which is a FIT target and upregulated under -Fe (Colangelo & Guerinot, 2004). FIT protein also interferes with Cu regulators (Cai et al., 2021; Perea-García et al., 2020). This observation raises the question whether the gene expression changes and leaf growth suppression can truly be attributed to -Fe or whether it could be a secondary consequence of elevated Cu. Nicotianamine is a metal chelator for Cu and Fe, and Cu homeostasis is equally important for reproduction than Fe (Sheng et al., 2021). -Fe was also reported to cause Zn and Mn accumulation in seedlings due to the elevated IRT1 activity leading to the uptake of other bivalent metals including Fe, Zn and Mn (Cointry & Vert, 2019; Dubeaux et al., 2018). Since we did not see this effect of Zn and Mn accumulation in -Fe in roots and rosette leaves at the reproductive stage, except for elevated Mn at MS in roots, it is possible that IRT1 function has a different relevance during reproduction or that mechanisms are in place to repress metal uptake and transport to rosette leaves targeting Mn and Zn, thus preventing over accumulation.

### Future perspectives

The results of this study indicate a clear and significant negative impact of Fe deficiency on rosette and young leaf growth dynamics, as well as the mineral partitioning in *Arabidopsis thaliana* during the reproductive stages. The findings underscore the crucial role of Fe in plant growth at reproductive stages and complex interactions and signaling between the various plant parts. This underlines that more attention should be paid to the roles of Fe homeostasis genes in the reproductive stages. Further analysis is thus required to elucidate the molecular mechanisms acting during reproduction. Our study was limited by two aspects. On one side, only wild type Col-0 plants were investigated. Future studies shall focus on natural variation and mutants of individual genes to undermine their roles in reproduction. On the other side, conclusions are based on gene expression and metal contents, excluding the physiological roles of the encoded transport proteins and enzymes. These components are subject of post-translational and protein control, and clearly mutants affected in such regulatory mechanisms can reveal the contributions during reproductive stages.

The question arises whether old leaves can be considered as Fe sources and whether young leaves are Fe sinks at +Fe (Hantzis et al., 2018; Nguyen et al., 2022; Swartz et al., 2023). The here-established growth system can be utilized to test such questions by performing Fe supply experiments in combination with mutants that cannot steer Fe physiology properly at the reproductive stage. In the future, it will be interesting to investigate the molecular mechanisms regulating potentially systemic Fe signals from source to sinks in the rosette, and link gene expression with leaf growth signaling under +Fe and -Fe, e.g. by utilizing SPL9 mutants or investigating the role of retrograde signaling pathway in leaf-to-leaf communication.

Finally, analyzing gene expressions at the reproductive phase is challenging, since variations in gene expression are relatively high and the whole plant physiology far more complex than in young seedlings. Whole transcriptome studies will offer better insight into the general physiology of old and young leaves and more complete investigation of Fe homeostasis genes in the future.

## Acknowledgements

The authors thank Sabine Metzger, University of Cologne, for technical support in the ICP-MS measurements. We acknowledge Rainer Franzen, Max Planck Institute for Plant Breeding Research Cologne, for help with SEM imaging of inflorescence shoot apices. We thank Elke Wieneke, Monique Eutebach and Florian Pollmeyer, all from Heinrich Heine University Düsseldorf, for excellent technical support. We thank Hans-Jörg Mai, Heinrich Heine University Düsseldorf, for his helpful suggestions regarding data visualization and manuscript figures. Chat GPT 4.0 has been used to edit and shorten text, correct and improve language. This work received funding by Deutsche Forschungsgemeinschaft (DFG, German Research Foundation) under Germanýs Excellence Strategy – EXC-2048/1 – project ID 390686111.

## Author contributions

Conceptualization (M.N., M.K., P.B.), Methodology (M.N., M.K., R.R.), Investigation (M.N., R.R.), Data Analysis (M.N., M.K., R.R., S.G.), Writing of Original Draft (M.N., M.K., P.B), Review & Editing (all authors), Supervision (M.K., P.B.), and Funding Acquisition (P.B.).

## Supplemental material

**Figure S1.**
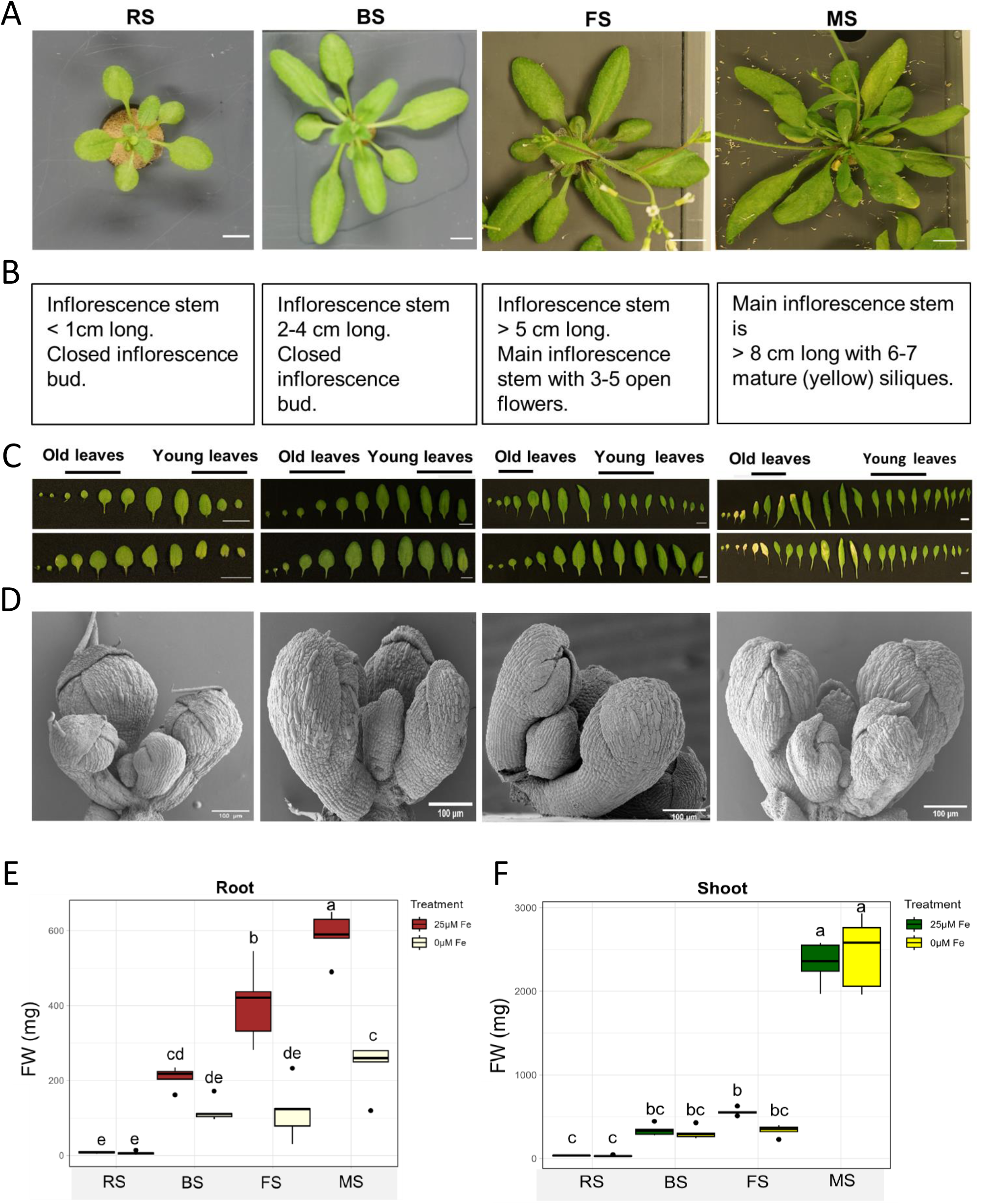
Materials collected from *Arabidopsis* Col-*0*. (A) Top view of plants with their rosette leaves at each stage, scale bar = 1cm. (B) Descriptions of the growth stages distinguishing phenotypes as described by (Boyes et al., 2001). (C) Old (left) and young (right) leaves collected from plants at the respective stages under +Fe (control) and –Fe after 3 days, scale bar = 1cm. (D) Inflorescence shoot apical meristem dissected from the main stem of plants at the rosette to mature stages., scale bar = 100 µm. (E-F) Root and shoot fresh weight of plants grown to RS, BS, FS, and MS and transferred to hydroponic media with +Fe or -Fe for 3 days. Data represents mean of three biological replicates (*n* = 15). Error bars show ± SD, one-way ANOVA and multiple comparison by Tukey test.

**Figure S2.**
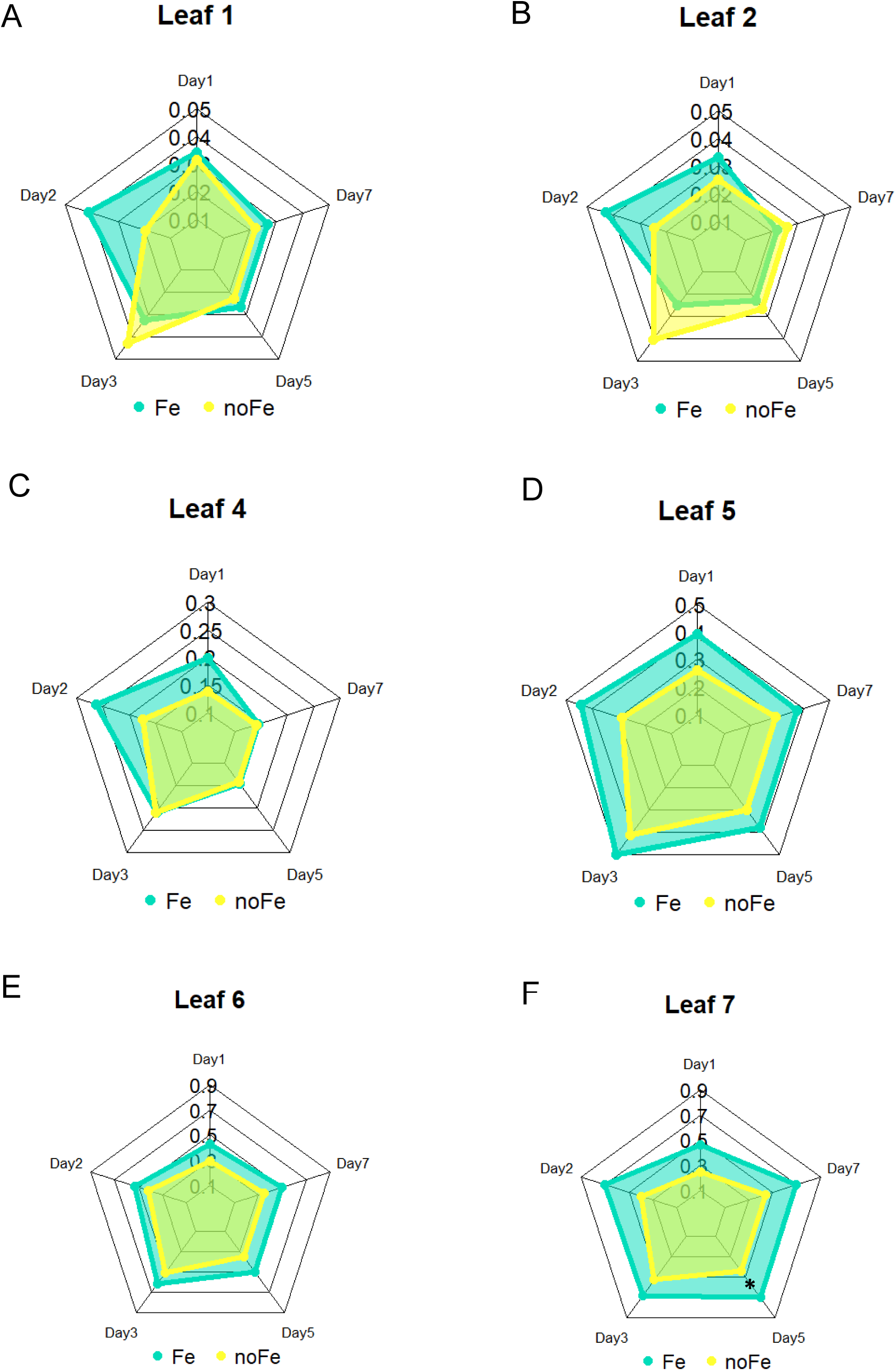
Individual sizes of leaves at RS after +Fe and -Fe. (A-F) Individual leaf areas of plants at RS after exposure to either Fe sufficient (+Fe) or deficient (-Fe) conditions from day 1 to day 7. Data represents average of biological replicates (*n*=3). Radial charts were plotted in R Studio using “fsmb” package. Asterisks show where leaf areas were statistically significantly different between +Fe and -Fe conditions, (p<0.05), one-way ANOVA and Tukey’s HSD test.

**Figure S3.**
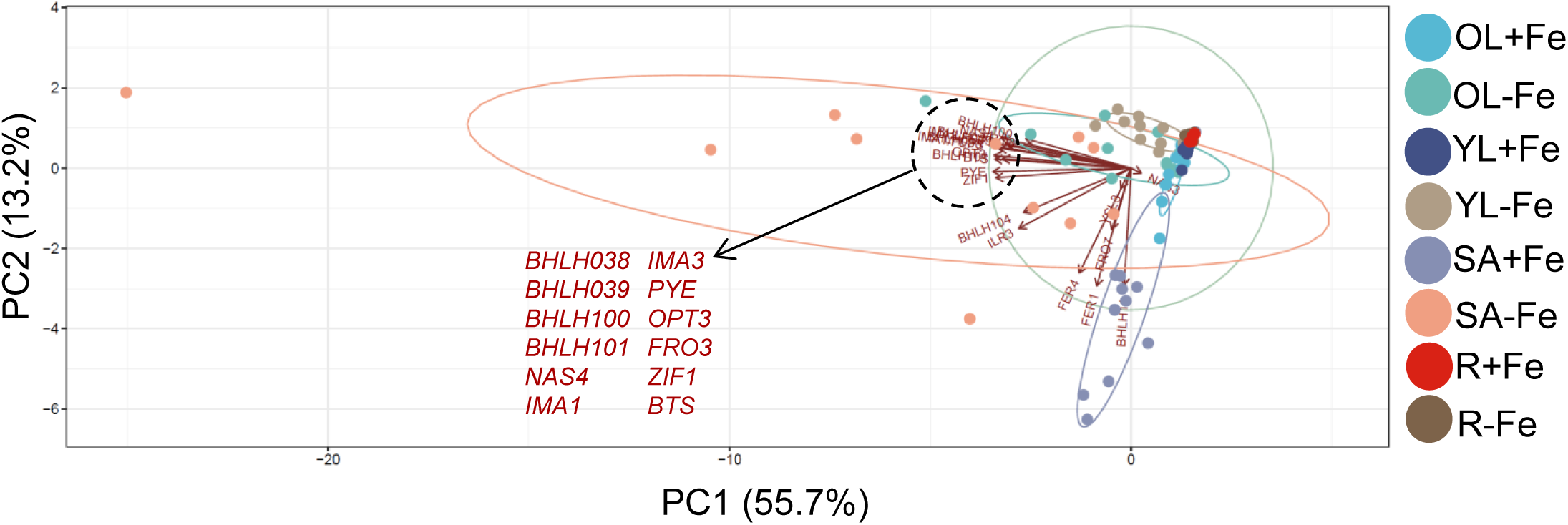
Principal component (PC) analysis with separation of old leaves, young leaves, shoot apex and roots according to gene expression at +Fe (control) or –Fe conditions at growth stages RS, BS, FS and MS. Samples are labeled as OL – old leaves, YL – young leaves, R – roots and SA – shoot apex. PC1 & PC2 separated samples according to growth stages and Fe condition. PCA analysis was computed using normalized absolute expression values of Fe-response genes.

**Figure S4.**
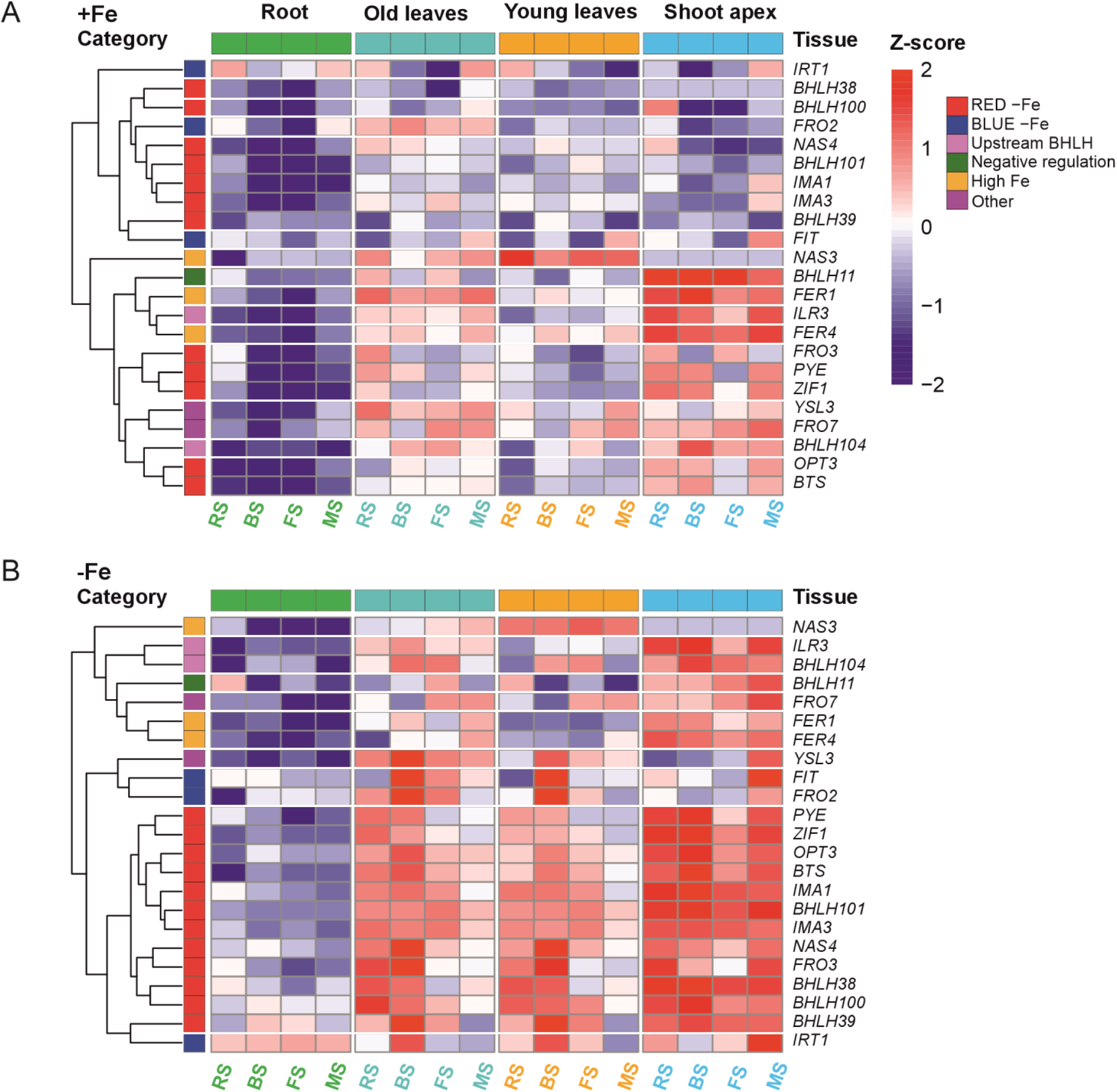
(A-B) Hierarchical clustering of the expression pattern of selected 23 Fe-deficiency responsive genes in roots, old leaves, young leaves, and inflorescence shoot apex at +Fe (control) and –Fe respectively, for 3 days during four stages; RS, BS, FS and MS. Gene category annotations are displayed on the left side and tissue-type annotations on the top. Gene upregulation is represented in red color and gene downregulation with blue colors, p < 0.05 (n = 3). The color scale ranges from -2 to 2 to represent the variation in gene expression.

**Figure S5.**
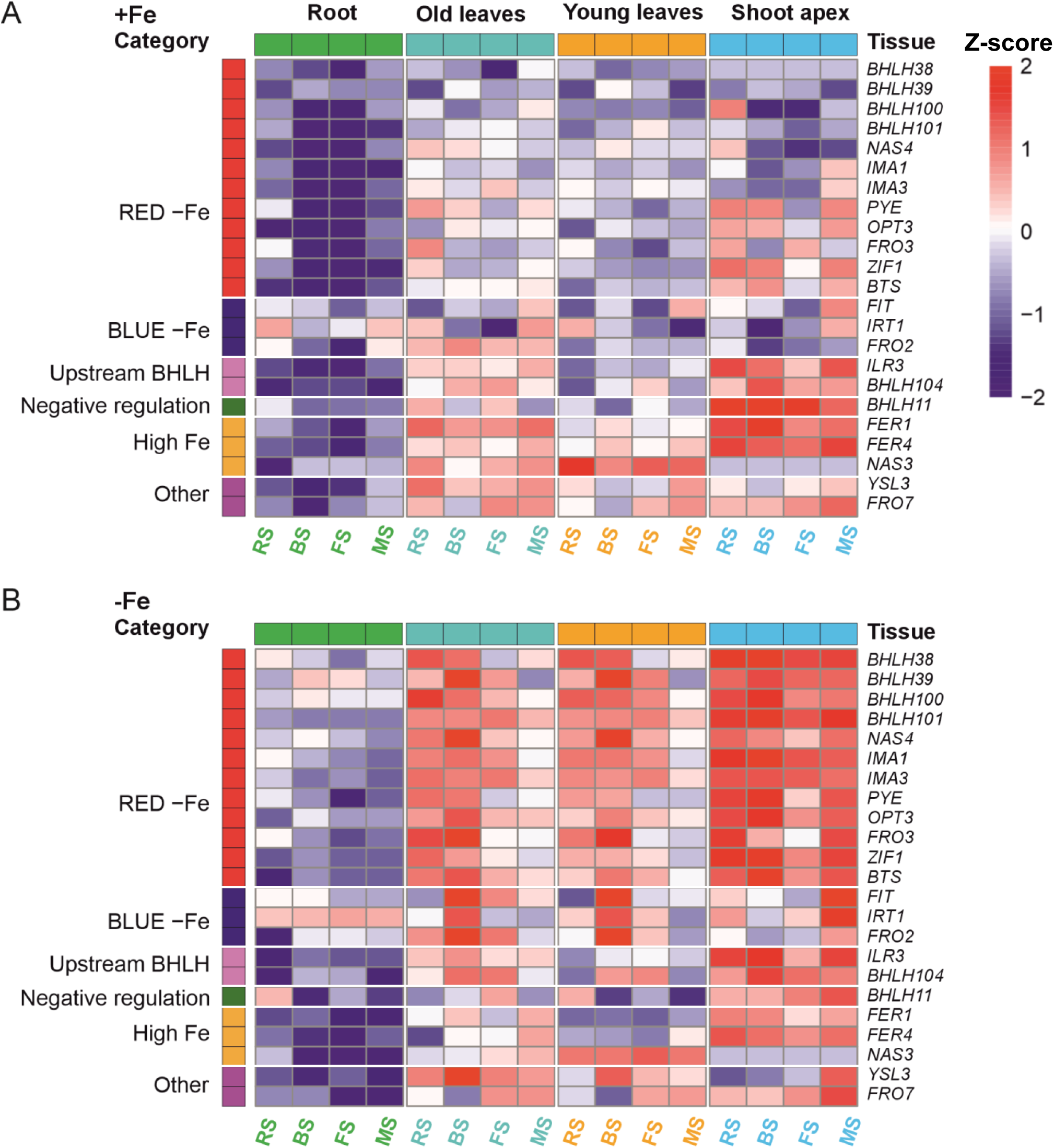
RT-qPCR analysis of Fe regulated genes across the stages. (A-B) Heatmaps showing the absolute expression profile of 23 Fe-response markers in roots, old leaves, young leaves, and inflorescence shoot apex at +Fe (control) and –Fe respectively, for 3 days during four stages; RS, BS, FS and MS. Functional gene category annotations are displayed on the left side of the heatmap, with tissue-type annotations displayed on the top. Gene upregulation is represented in red color and gene downregulation with blue colors, p < 0.05 (n = 3). The color scale ranges from -2 to 2 to represent the variation in gene expression.

**Figure S6.**
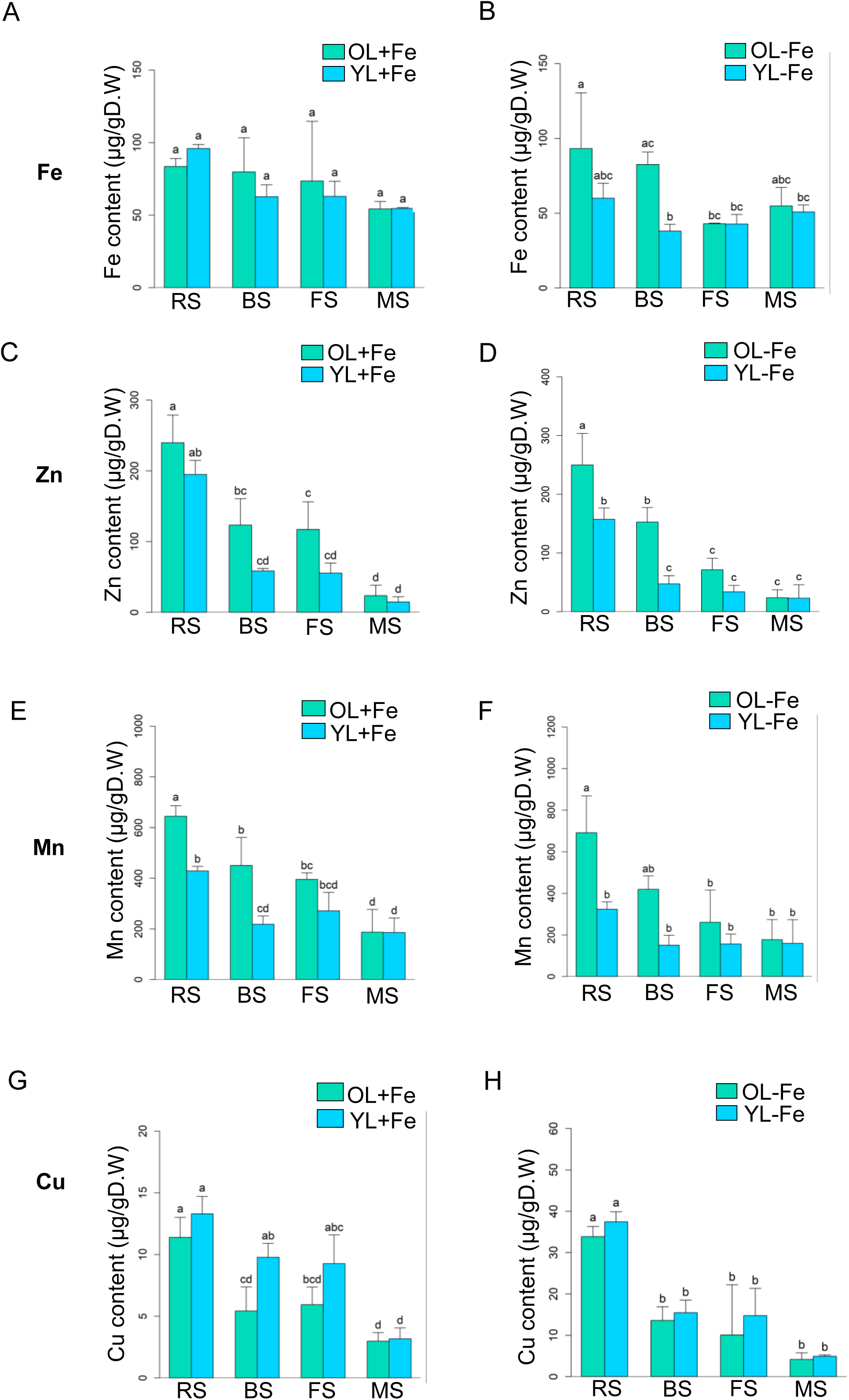
Comparison of mineral contents in old (OL) and young (YL) leaves at +Fe and –Fe growth conditions. Fe (A-B), Zn (C-D), Mn (E-F), and Cu (G-H) content in old leaves and young leaves at +Fe or –Fe growth conditions at the growth stages; RS, BS, FS and MS. Plants were exposed to 3 days of +Fe or –Fe before sample collection. Data represents mean of three biological replicates (*n* = 15). Error bars represent ± SD. Letters on top of each bar show level of significance according to one-way ANOVA and Tukey’s HSD test, p < 0.05.

**Figure S7.**
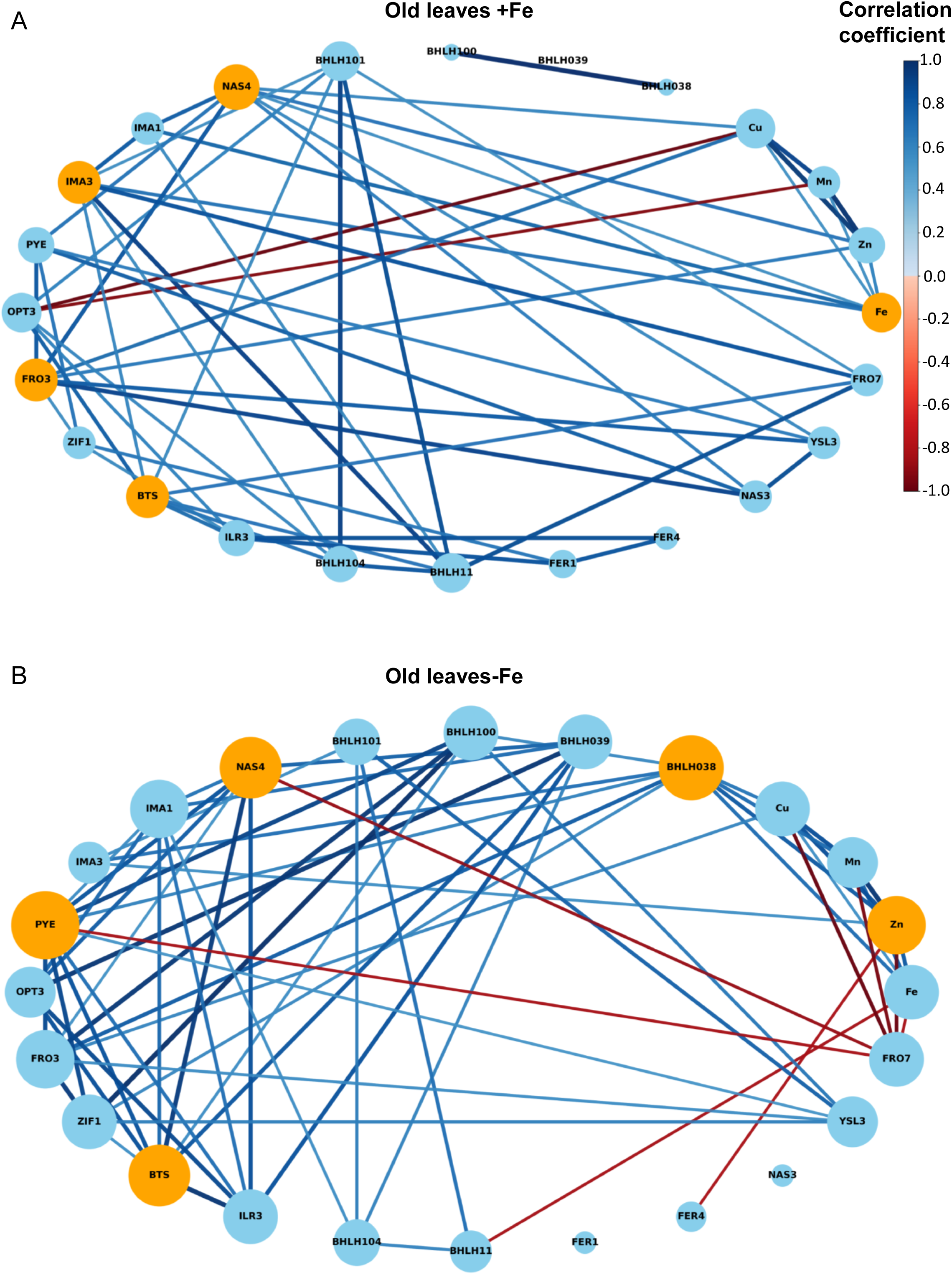
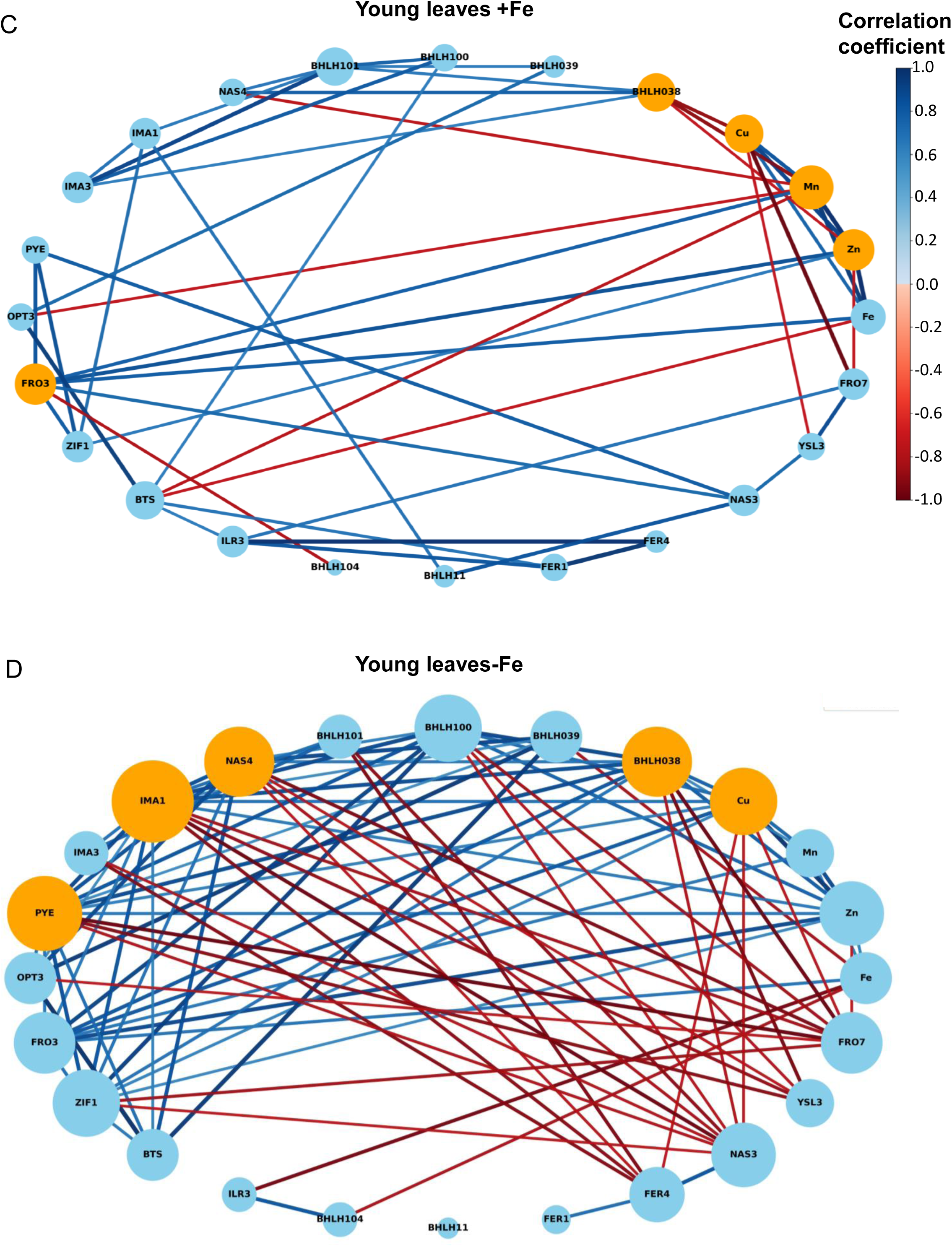
Network analysis of correlations between genes and minerals Fe, Mn, Zn and Cu in old (A & B) and young leaves (C & D) at +Fe and –Fe conditions, respectively. Node sizes represent the degree centrality of each node, with larger nodes indicating genes or minerals with higher degree centrality while smaller nodes show genes with lower degree centrality. Orange nodes highlight most connected genes. Edge colors represent correlation, positive correlations are shown in blue and red for negative correlations. Edge thickness represent the strength of the correlation between connected nodes. Correlations that were above 0.5 are displayed, p value<0.05. Scale ranges from 1 to -1 to account for variation in correlation.

**Table S1.**
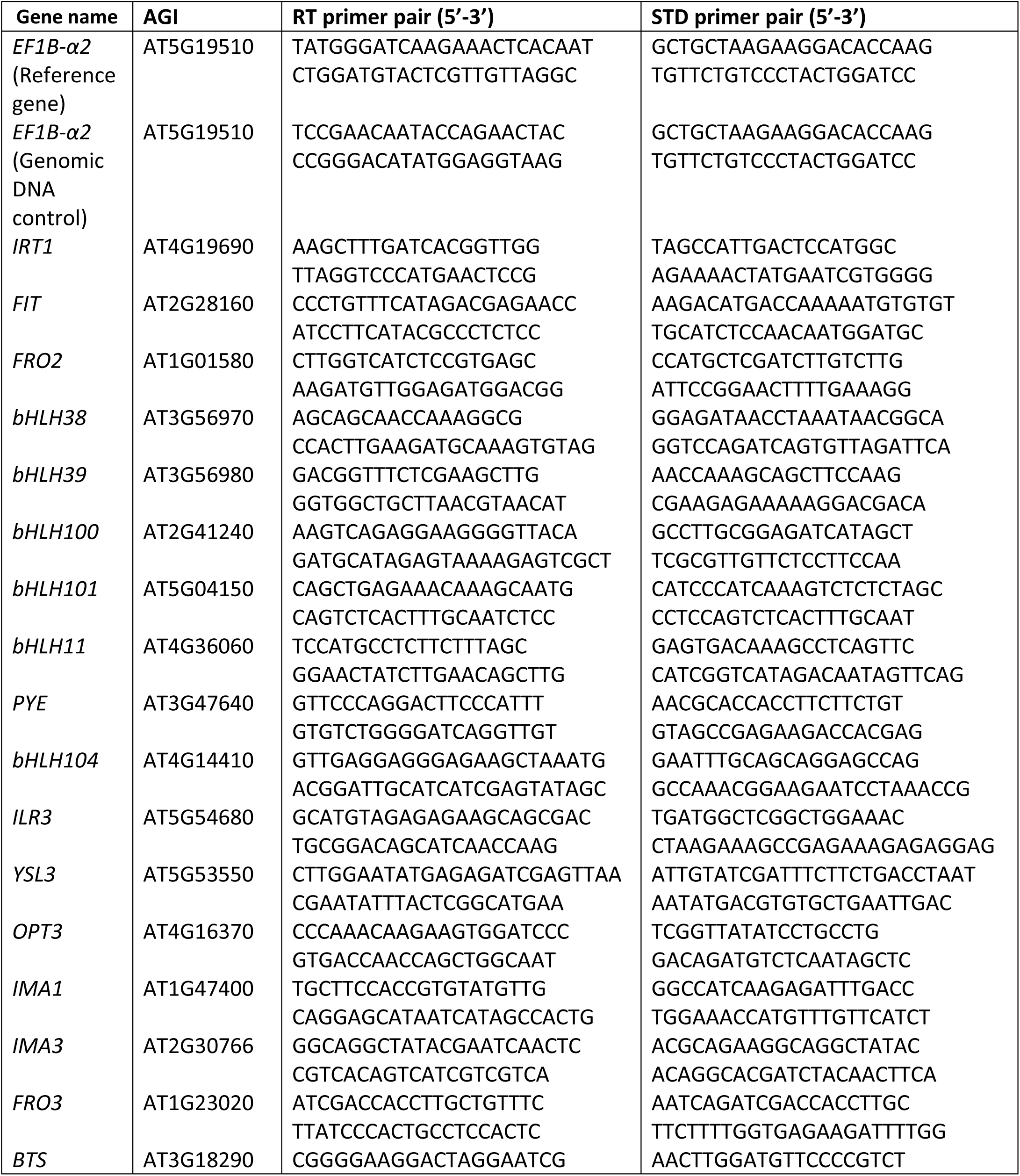
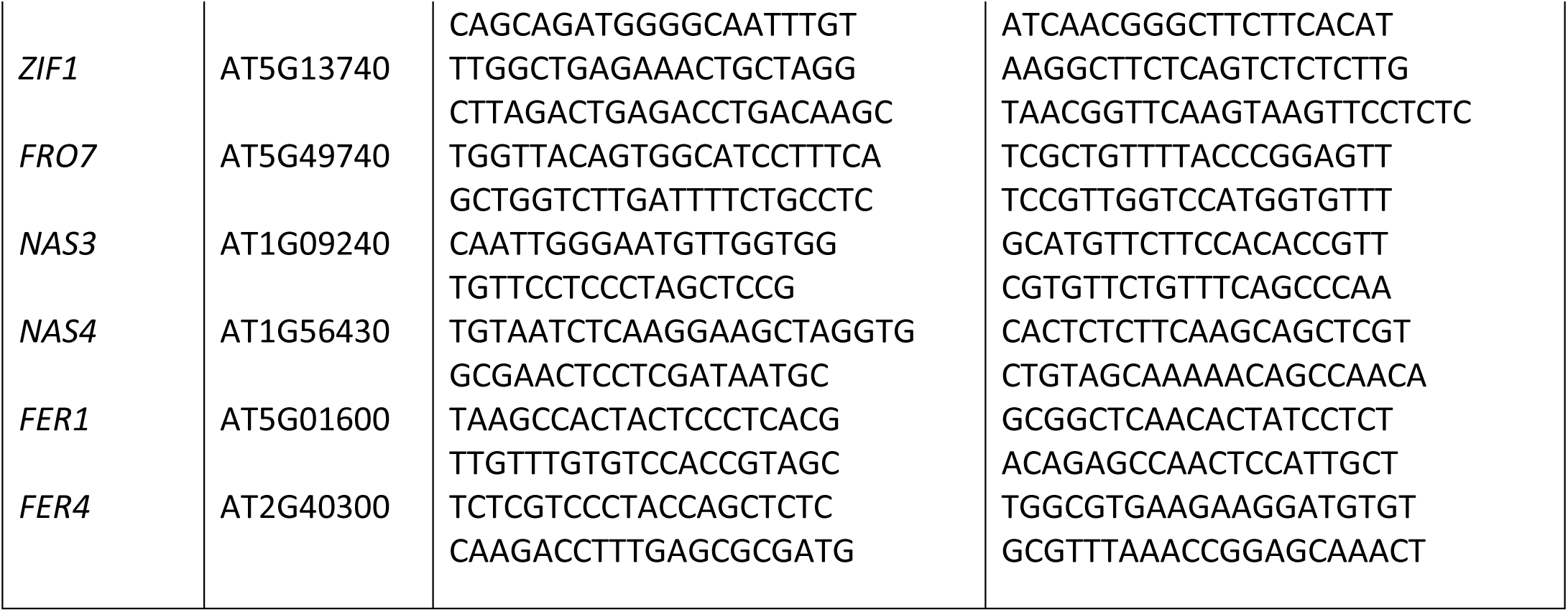
Primers for qPCR. List of primers used for gene expression analysis by reverse transcription-qPCR. Primers were designed as previously described in (Ngigi & Bauer, 2023). Mass standard (STD) and real time qPCR (RT) primers annealed to the indicated genes. Mass standard PCR products were used for absolute quantification of transcript abundances of the respective genes.

## Notes

### Competing Interest Statement

The authors have declared no competing interest.

### Summary of Updates

- added references into introduction - revised text according to reviewer comments - reanalyzed further data and provided respective figures and supplemental figures - provided statistical analysis for leaf areas (in radial charts)

## References

Akmakjian, G. Z., Riaz, N., & Guerinot, M. L. (2021). Photoprotection during iron deficiency is mediated by the bHLH transcription factors PYE and ILR3. Proceedings of the National Academy of Sciences, 118(40). 10.1073/PNAS.2024918118

Andriankaja, M., Dhondt, S., De Bodt, S., Vanhaeren, H., Coppens, F., De Milde, L., Mühlenbock, P., Skirycz, A., Gonzalez, N., Beemster, G. T. S., & Inzé, D. (2012). Exit from Proliferation during Leaf Development in *Arabidopsis thaliana*: A Not-So-Gradual Process. Developmental Cell, 22(1), 64–78. 10.1016/j.devcel.2011.11.011

Andriankaja, M. E., Danisman, S., Mignolet-Spruyt, L. F., Claeys, H., Kochanke, I., Vermeersch, M., De Milde, L., De Bodt, S., Storme, V., Skirycz, A., Maurer, F., Bauer, P., Mühlenbock, P., Van Breusegem, F., Angenent, G. C., Immink, R. G. H., & Inzé, D. (2014). Transcriptional coordination between leaf cell differentiation and chloroplast development established by TCP20 and the subgroup Ib bHLH transcription factors. Plant Molecular Biology, 85(3), 233–245. 10.1007/s11103-014-0180-2

Balparda, M., Armas, A. M., Estavillo, G. M., Roschzttardtz, H., Pagani, M. A., & Gomez-Casati, D. F. (2020). The PAP/SAL1 retrograde signaling pathway is involved in iron homeostasis. Plant Molecular Biology, 102(3), Article 3. 10.1007/s11103-019-00950-7

Balparda, M., Armas, A. M., Gomez-Casati, D. F., & Pagani, M. A. (2021). PAP/SAL1 retrograde signaling pathway modulates iron deficiency response in alkaline soils. Plant Science, 304, 110808. 10.1016/j.plantsci.2020.110808

Bastow, E. L., Garcia de la Torre, V. S., Maclean, A. E., Green, R. T., Merlot, S., Thomine, S., & Balk, J. (2018). Vacuolar iron stores gated by NRAMP3 and NRAMP4 are the primary source of iron in germinating seeds. Plant Physiology, 177(3), 1267–1276. 10.1104/pp.18.00478

Bäurle, I., & Dean, C. (2006). The Timing of Developmental Transitions in Plants. Cell. https://www.cell.com/cell/abstract/S0092-8674(06)00571-X

Bernal, M., Casero, D., Singh, V., Wilson, G. T., Grande, A., Yang, H., Dodani, S. C., Pellegrini, M., Huijser, P., Connolly, E. L., Merchant, S. S., & Krämer, U. (2012). Transcriptome Sequencing Identifies SPL7-Regulated Copper Acquisition Genes FRO4/FRO5 and the Copper Dependence of Iron Homeostasis in Arabidopsis. The Plant Cell, 24(2), 738–761. 10.1105/tpc.111.090431

Boyes, D. C., Zayed, A. M., Ascenzi, R., Mccaskill, A. J., Hoffman, N. E., Davis, K. R., & Görlach, J. (2001). Growth Stage-Based Phenotypic Analysis of Arabidopsis: A Model for High Throughput Functional Genomics in Plants. The Plant Cell, 13, 1499–1510.

Briat, J. F., Curie, C., & Gaymard, F. (2007). Iron utilization and metabolism in plants. Current Opinion in Plant Biology, 10(3), 276–282. 10.1016/J.PBI.2007.04.003

Briat, J.-F., Ravet, K., Arnaud, N., Duc, C., Boucherez, J., Touraine, B., Cellier, F., & Gaymard, F. (2010). New insights into ferritin synthesis and function highlight a link between iron homeostasis and oxidative stress in plants. Annals of Botany, 105(5), 811–822. 10.1093/aob/mcp128

Buchanan-Wollaston, V. (1997). The molecular biology of leaf senescence. Journal of Experimental Botany, 48(2), 181–199. 10.1093/jxb/48.2.181

Cai, Y., Li, Y., & Liang, G. (2021). FIT and bHLH Ib transcription factors modulate iron and copper crosstalk in Arabidopsis. Plant, Cell & Environment, 44(5), 1679–1691. 10.1111/pce.14000

Carrió-Seguí, À., Romero, P., Curie, C., Mari, S., & Peñarrubia, L. (2019). Copper transporter COPT5 participates in the crosstalk between vacuolar copper and iron pools mobilisation. Scientific Reports, 9(1), Article 1. 10.1038/s41598-018-38005-4

Chia, J.-C., Yan, J., Ishka, M. R., Faulkner, M. M., Simons, E., Huang, R., Smieska, L., Woll, A., Tappero, R., Kiss, A., Jiao, C., Fei, Z., Kochian, L. V., Walker, E., Piñeros, M., & Vatamaniuk, O. K. (2023). Loss of OPT3 function decreases phloem copper levels and impairs crosstalk between copper and iron homeostasis and shoot-to-root signaling in *Arabidopsis thaliana*. *The Plant Cell*, koad053. 10.1093/plcell/koad053

Chu, H.-H., Chiecko, J., Punshon, T., Lanzirotti, A., Lahner, B., Salt, D. E., & Walker, E. L. (2010). Successful Reproduction Requires the Function of Arabidopsis YELLOW STRIPE-LIKE1 and YELLOW STRIPE-LIKE3 Metal-Nicotianamine Transporters in Both Vegetative and Reproductive Structures 1[W][OA]. Plant Physiology Ò, 154, 197–210. 10.1104/pp.110.159103

Cointry, V., & Vert, G. (2019). The bifunctional transporter-receptor IRT1 at the heart of metal sensing and signalling. New Phytologist, 223(3), 1173–1178. 10.1111/nph.15826

Colangelo, E. P., & Guerinot, M. L. (2004). The Essential Basic Helix-Loop-Helix Protein FIT1 Is Required for the Iron Deficiency Response. The Plant Cell, 16(12), 3400–3412. 10.1105/tpc.104.024315

Costa, M. M. R., Yang, S., Critchley, J., Feng, X., Wilson, Y., Langlade, N., Copsey, L., & Hudson, A. (2012). The genetic basis for natural variation in heteroblasty in Antirrhinum. New Phytologist, 196(4), 1251–1259. 10.1111/j.1469-8137.2012.04347.x

Dubeaux, G., Neveu, J., Zelazny, E., & Vert, G. (2018). Metal Sensing by the IRT1 Transporter-Receptor Orchestrates Its Own Degradation and Plant Metal Nutrition. Molecular Cell, 69(6), 953–964.e5. 10.1016/j.molcel.2018.02.009

Hantzis, L. J., Kroh, G. E., Jahn, C. E., Cantrell, M., Peers, G., Pilon, M., & Raveta, K. (2018). A Program for Iron Economy during Deficiency Targets Specific Fe Proteins. Plant Physiology, 176(1), 596– 610. 10.1104/PP.17.01497

Haydon, M. J., & Cobbett, C. S. (2007). A Novel Major Facilitator Superfamily Protein at the Tonoplast Influences Zinc Tolerance and Accumulation in Arabidopsis. Plant Physiology, 143(4), 1705– 1719. 10.1104/pp.106.092015

Haydon, M. J., Kawachi, M., Wirtz, M., Hillmer, S., Diger Hell, R., & Krämer, U. (2012). Vacuolar nicotianamine has critical and distinct roles under iron deficiency and for zinc sequestration in Arabidopsis. The Plant Cell. 10.1105/tpc.111.095042

Himelblau, E., & Amasino, R. M. (2001). Nutrients mobilized from leaves of Arabidopsis thaliana during leaf senescence. Journal of Plant Physiology, 158(10), 1317–1323. 10.1078/0176-1617-00608

Ivanov, R., Brumbarova, T., & Bauer, P. (2012a). Fitting into the Harsh Reality: Regulation of Iron-deficiency Responses in Dicotyledonous Plants. Molecular Plant, 5(1), 27–42. 10.1093/mp/ssr065

Ivanov, R., Brumbarova, T., & Bauer, P. (2012b). Fitting into the harsh reality: Regulation of iron-deficiency responses in dicotyledonous plants. Molecular Plant, 5(1), 27–42. 10.1093/mp/ssr065

Jakoby, M., Wang, H. Y., Reidt, W., Weisshaar, B., & Bauer, P. (2004). FRU (BHLH029) is required for induction of iron mobilization genes in Arabidopsis thaliana. FEBS Letters, 577(3), 528–534. 10.1016/j.febslet.2004.10.062

Jeong, J., & Guerinot, M. L. (2009). Homing in on iron homeostasis in plants. Trends in Plant Science, 14(5), 280–285. 10.1016/j.tplants.2009.02.006

Kastoori Ramamurthy, R., Xiang, Q., Hsieh, E.-J., Liu, K., Zhang, C., & Waters, B. M. (2018). New aspects of iron–copper crosstalk uncovered by transcriptomic characterization of Col-0 and the copper uptake mutant spl7 in Arabidopsis thaliana†. Metallomics, 10(12), 1824–1840. 10.1039/c8mt00287h

Khan, M. A., Castro-Guerrero, N. A., McInturf, S. A., Nguyen, N. T., Dame, A. N., Wang, J., Bindbeutel, R. K., Joshi, T., Jurisson, S. S., Nusinow, D. A., & Mendoza-Cozatl, D. G. (2018). Changes in iron availability in Arabidopsis are rapidly sensed in the leaf vasculature and impaired sensing leads to opposite transcriptional programs in leaves and roots. *Plant*, Cell & Environment, 41(10), 2263–2276. 10.1111/PCE.13192

Kobayashi, T., Nagano, A. J., & Nishizawa, N. K. (2021). Iron deficiency-inducible peptide-coding genes OsIMA1 and OsIMA2 positively regulate a major pathway of iron uptake and translocation in rice. Journal of Experimental Botany, 72(6), 2196–2211. 10.1093/jxb/eraa546

Kumar, R. K., Chu, H. H., Abundis, C., Vasques, K., Rodriguez, D. C., Chia, J. C., Huang, R., Vatamaniuk, O. K., & Walker, E. L. (2017). Iron-Nicotianamine Transporters Are Required for Proper Long Distance Iron Signaling. Plant Physiology, 175(3), 1254–1268. 10.1104/PP.17.00821

Lanquar, V., Lelièvre, F., Bolte, S., Hamès, C., Alcon, C., Neumann, D., Vansuyt, G., Curie, C., Schröder, A., Krämer, U., Barbier-Brygoo, H., & Thomine, S. (2005). Mobilization of vacuolar iron by AtNRAMP3 and AtNRAMP4 is essential for seed germination on low iron. The EMBO Journal, 24(23), 4041. 10.1038/SJ.EMBOJ.7600864

Le, C. T. T., Brumbarova, T., Ivanov, R., Stoof, C., Weber, E., Mohrbacher, J., Fink-Straube, C., & Bauer, P. (2016). ZINC FINGER OF ARABIDOPSIS THALIANA12 (ZAT12) Interacts with FER-LIKE IRON DEFICIENCY-INDUCED TRANSCRIPTION FACTOR (FIT) Linking Iron Deficiency and Oxidative Stress Responses. Plant Physiology, 170(1), 540–557. 10.1104/pp.15.01589

Lehmann, F., & Hardtke, C. S. (2016). Secondary growth of the Arabidopsis hypocotyl—Vascular development in 4 dimensions. Current Opinion in Plant Biology, 29, 9–15. 10.1016/j.pbi.2015.10.011

Lemaître, T., Gaufichon, L., Boutet-Mercey, S., Christ, A., & Masclaux-Daubresse, C. (2008). Enzymatic and Metabolic Diagnostic of Nitrogen Deficiency in Arabidopsis thaliana Wassileskija Accession. Plant and Cell Physiology, 49(7), 1056–1065. 10.1093/pcp/pcn081

Li, X., Zhang, H., Ai, Q., Liang, G., & Yu, D. (2016). Two bHLH transcription factors, bHLH34 and bHLH104, regulate iron homeostasis in arabidopsis Thaliana. Plant Physiology, 170(4), 2478–2493. 10.1104/pp.15.01827

Li, X.-M., Jenke, H., Strauss, S., Bazakos, C., Mosca, G., Lymbouridou, R., Kierzkowski, D., Neumann, U., Naik, P., Huijser, P., Laurent, S., Smith, R. S., Runions, A., & Tsiantis, M. (2024a). Cell-cycle-linked growth reprogramming encodes developmental time into leaf morphogenesis. Current Biology, 34(3), 541–556.e15. 10.1016/j.cub.2023.12.050

Li, X.-M., Jenke, H., Strauss, S., Bazakos, C., Mosca, G., Lymbouridou, R., Kierzkowski, D., Neumann, U., Naik, P., Huijser, P., Laurent, S., Smith, R. S., Runions, A., & Tsiantis, M. (2024b). Cell-cycle-linked growth reprogramming encodes developmental time into leaf morphogenesis. Current Biology, 34(3), 541–556.e15. 10.1016/j.cub.2023.12.050

Lichtblau, D. M., Schwarz, B., Baby, D., Endres, C., Sieberg, C., & Bauer, P. (2022). The Iron Deficiency-Regulated Small Protein Effector FEP3/IRON MAN1 Modulates Interaction of BRUTUS-LIKE1 With bHLH Subgroup IVc and POPEYE Transcription Factors. Frontiers in Plant Science, 13(June), 1–22. 10.3389/fpls.2022.930049

Long, T. A., Tsukagoshi, H., Busch, W., Lahner, B., Salt, D. E., & Benfey, P. N. (2010). The bHLH Transcription Factor POPEYE Regulates Response to Iron Deficiency in Arabidopsis Roots. The Plant Cell, 22(7), 2219–2236. 10.1105/tpc.110.074096

Mai, H.-J., Baby, D., & Bauer, P. (2023). Black sheep, dark horses, and colorful dogs: A review on the current state of the Gene Ontology with respect to iron homeostasis in Arabidopsis thaliana. Frontiers in Plant Science. 10.3389/fpls.2023.1204723

Mai, H.-J., Pateyron, S., & Bauer, P. (2016). Iron homeostasis in Arabidopsis thaliana: Transcriptomic analyses reveal novel FIT-regulated genes, iron deficiency marker genes and functional gene networks. BMC Plant Biology, 16(1), 211. 10.1186/s12870-016-0899-9

Meiser, J., Lingam, S., & Bauer, P. (2011). Posttranslational Regulation of the Iron Deficiency Basic Helix-Loop-Helix Transcription Factor FIT Is Affected by Iron and Nitric Oxide. Plant Physiology, 157(4), 2154–2166. 10.1104/pp.111.183285

Mendoza-Cózatl, D. G., Xie, Q., Akmakjian, G. Z., Jobe, T. O., Patel, A., Stacey, M. G., Song, L., Demoin, D. W., Jurisson, S. S., Stacey, G., & Schroeder, J. I. (2014). OPT3 Is a Component of the Iron-Signaling Network between Leaves and Roots and Misregulation of OPT3 Leads to an Over-Accumulation of Cadmium in Seeds. Molecular Plant, 7(9), 1455. 10.1093/MP/SSU067

Muhammad, D., Clark, N. M., Haque, S., Williams, C. M., Sozzani, R., & Long, T. A. (2022). POPEYE intercellular localization mediates cell-specific iron deficiency responses. Plant Physiology, 190(3), 2017–2032. 10.1093/plphys/kiac357

Mukherjee, I., Campbell, N. H., Ash, J. S., & Connolly, E. L. (2006). Expression profiling of the Arabidopsis ferric chelate reductase (FRO) gene family reveals differential regulation by iron and copper. Planta, 223(6), 1178–1190. 10.1007/s00425-005-0165-0

Nam, H.-I., Shahzad, Z., Dorone, Y., Clowez, S., Zhao, K., Bouain, N., Lay-Pruitt, K. S., Cho, H., Rhee, S. Y., & Rouached, H. (2021). Interdependent iron and phosphorus availability controls photosynthesis through retrograde signaling. Nature Communications, 12(1), Article 1. 10.1038/s41467-021-27548-2

Naranjo-Arcos, M. A., Maurer, F., Meiser, J., Pateyron, S., Fink-Straube, C., & Bauer, P. (2017). Dissection of iron signaling and iron accumulation by overexpression of subgroup Ib bHLH039 protein. Scientific Reports 2017 7:1, 7(1), 1–12. 10.1038/s41598-017-11171-7

Ngigi, M. N., & Bauer, P. (2023). High-Throughput Plant Gene Expression Analysis by 384-Format Reverse Transcription-Quantitative PCR for Investigating Plant Iron Homeostasis. In J. Jeong (Ed.), Plant Iron Homeostasis: Methods and Protocols (pp. 1–22). Springer US. 10.1007/978-1-0716-3183-6_1

Nguyen, N. T., Khan, M. A., Castro–Guerrero, N. A., Chia, J.-C., Vatamaniuk, O. K., Mari, S., Jurisson, S. S., & Mendoza-Cozatl, D. G. (2022). Iron Availability within the Leaf Vasculature Determines the Magnitude of Iron Deficiency Responses in Source and Sink Tissues in Arabidopsis. Plant and Cell Physiology. 10.1093/PCP/PCAC046

Nguyen, N. T., McInturf, S. A., & Mendoza-Cózatl, D. G. (2016). Hydroponics: A Versatile System to Study Nutrient Allocation and Plant Responses to Nutrient Availability and Exposure to Toxic Elements. Journal of Visualized Experiments : JoVE, 113, 54317. 10.3791/54317

Nooden, L. D., Hillsberg, J. W., & Schneider, M. J. (1996). Induction of leaf senescence in Arabidopsis thaliana by long days through a light-dosage effect. Physiologia Plantarum, 96(3), 491–495. 10.1111/j.1399-3054.1996.tb00463.x

Perea-García, A., Andrés-Bordería, A., Vera-Sirera, F., Pérez-Amador, M. A., Puig, S., & Peñarrubia, L. (2020). Deregulated High Affinity Copper Transport Alters Iron Homeostasis in Arabidopsis. Frontiers in Plant Science. 10.3389/fpls.2020.01106

Pottier, M., Dumont, J., Masclaux-Daubresse, C., & Thomine, S. (2019a). Autophagy is essential for optimal translocation of iron to seeds in Arabidopsis. Journal of Experimental Botany, 70(3), 859– 869. 10.1093/jxb/ery388

Pottier, M., Dumont, J., Masclaux-Daubresse, C., & Thomine, S. (2019b). Autophagy is essential for optimal translocation of iron to seeds in Arabidopsis. Journal of Experimental Botany, 70(3), 859– 869. 10.1093/jxb/ery388

Pu, M. N., & Liang, G. (2023). The transcription factor POPEYE negatively regulates the expression of bHLH Ib genes to maintain iron homeostasis. Journal of Experimental Botany, 74(8), 2754– 2767. 10.1093/jxb/erad057

Rai, S., Singh, P. K., Mankotia, S., Swain, J., & Satbhai, S. B. (2021). Iron homeostasis in plants and its crosstalk with copper, zinc, and manganese. Plant Stress, 1, 100008. 10.1016/j.stress.2021.100008

Rodríguez-Celma, J., Connorton, J. M., Kruse, I., Green, R. T., Franceschetti, M., Chen, Y. T., Cui, Y., Ling, H. Q., Yeh, K. C., & Balk, J. (2019). Arabidopsis BRUTUS-LIKE E3 ligases negatively regulate iron uptake by targeting transcription factor FIT for recycling. Proceedings of the National Academy of Sciences of the United States of America, 116(35), 17584–17591. 10.1073/pnas.1907971116

Roschzttardtz, H., Conéjéro, G., Divol, F., Alcon, C., Verdeil, J.-L., Curie, C., & Mari, S. (2013). New insights into Fe localization in plant tissues. Frontiers in Plant Science. https://www.frontiersin.org/journals/plant-science/articles/10.3389/fpls.2013.00350/full

Sági-Kazár, M., Solymosi, K., & Solti, Á. (2022). Iron in leaves: Chemical forms, signalling, and in-cell distribution. Journal of Experimental Botany, 73(6), 1717–1734. 10.1093/jxb/erac030

Saini, K., Dwivedi, A., & Ranjan, A. (2022). High temperature restricts cell division and leaf size by coordination of PIF4 and TCP4 transcription factors. Plant Physiology, 190(4), 2380–2397. 10.1093/plphys/kiac345

Schuler, M., Rellán-Álvarez, R., Fink-Straube, C., Abadía, J., & Bauera, P. (2012). Nicotianamine functions in the phloem-based transport of iron to sink organs, in pollen development and pollen tube growth in Arabidopsis. Plant Cell, 24(6), 2380–2400. 10.1105/tpc.112.099077

Schwarz, B., & Bauer, P. (2020). FIT, a regulatory hub for iron deficiency and stress signaling in roots, and FIT-dependent and -independent gene signatures. Journal of Experimental Botany, 71(5), 1694–1705. 10.1093/jxb/eraa012

Selote, D., Samira, R., Matthiadis, A., Gillikin, J. W., & Long, T. A. (2015). Iron-binding e3 ligase mediates iron response in plants by targeting basic helix-loop-helix transcription factors1[open]. Plant Physiology, 167(1), 273–286. 10.1104/pp.114.250837

Sheng, H., Jiang, Y., Rahmati, M., Chia, J.-C., Dokuchayeva, T., Kavulych, Y., Zavodna, T.-O., Mendoza, P. N., Huang, R., Smieshka, L. M., Miller, J., Woll, A. R., Terek, O. I., Romanyuk, N. D., Piñeros, M., Zhou, Y., & Vatamaniuk, O. K. (2021). YSL3-mediated copper distribution is required for fertility, seed size and protein accumulation in Brachypodium. Plant Physiology, 186(1), 655–676. 10.1093/plphys/kiab054

Sperotto, R. A., Vasconcelos, M. W., Grusak, M. A., & Fett, J. P. (2012). Effects of different Fe supplies on mineral partitioning and remobilization during the reproductive development of rice (Oryza sativa L.). Rice, 5(1), 27. 10.1186/1939-8433-5-27

Spielmann, J., Fanara, S., Cotelle, V., & Vert, G. (2023). Multilayered regulation of iron homeostasis in Arabidopsis. Frontiers in Plant Science, 14. https://www.frontiersin.org/articles/10.3389/fpls.2023.1250588

Stacey, M. G., Patel, A., McClain, W. E., Mathieu, M., Remley, M., Rogers, E. E., Gassmann, W., Blevins, D. G., & Stacey, G. (2008). The Arabidopsis AtOPT3 Protein Functions in Metal Homeostasis and Movement of Iron to Developing Seeds. Plant Physiology, 146(2), 589. 10.1104/PP.107.108183

Swartz, L. G., Liu, S., Dahlquist, D., Kramer, S. T., Walter, E. S., McInturf, S. A., Bucksch, A., & Mendoza-Cózatl, D. G. (2023). OPEN leaf: An open-source cloud-based phenotyping system for tracking dynamic changes at leaf-specific resolution in Arabidopsis. The Plant Journal, 116(6), 1600–1616. 10.1111/tpj.16449

Tanabe, N., Noshi, M., Mori, D., Nozawa, K., Tamoi, M., & Shigeoka, S. (2018). The basic helix-loop-helix transcription factor, bHLH11 functions in the iron-uptake system in Arabidopsis thaliana. Journal of Plant Research, 132(1), Article 1. 10.1007/s10265-018-1068-z

Tanabe, N., Noshi, M., Mori, D., Nozawa, K., Tamoi, M., & Shigeoka, S. (2019). The basic helix-loop-helix transcription factor, bHLH11 functions in the iron-uptake system in Arabidopsis thaliana. Journal of Plant Research, 132(1), 93–105. 10.1007/s10265-018-1068-z

Thieme, C. J., Rojas-Triana, M., Stecyk, E., Schudoma, C., Zhang, W., Yang, L., Miñambres, M., Walther, D., Schulze, W. X., Paz-Ares, J., Scheible, W.-R., & Kragler, F. (2015). Endogenous Arabidopsis messenger RNAs transported to distant tissues. Nature Plants, 1(4), 1–9. 10.1038/nplants.2015.25

Tissot, N., Robe, K., Gao, F., Grant-Grant, S., Boucherez, J., Bellegarde, F., Maghiaoui, A., Marcelin, R., Izquierdo, E., Benhamed, M., Martin, A., Vignols, F., Roschzttardtz, H., Gaymard, F., Briat, J. F., & Dubos, C. (2019). Transcriptional integration of the responses to iron availability in Arabidopsis by the bHLH factor ILR3. New Phytologist, 223(3), 1433–1446. 10.1111/nph.15753

Trofimov, K., Ivanov, R., Eutebach, M., Acaroglu, B., Mohr, I., Bauer, P., & Brumbarova, T. (2019). Mobility and localization of the iron deficiency-induced transcription factor bHLH039 change in the presence of FIT. Plant Direct, 3(12), e00190. 10.1002/pld3.190

Tsukaya, H. (2005). Leaf shape: Genetic controls and environmental factors. The International Journal of Developmental Biology, 49(5–6), 547–555. 10.1387/ijdb.041921ht

Tsukaya, H., Shoda, K., Kim, G. T., & Uchimiya, H. (2000). Heteroblasty in Arabidopsis thaliana (L.) Heynh. Planta, 210(4), 536–542. 10.1007/s004250050042

Vert, G., Grotz, N., Dédaldéchamp, F., Gaymard, F., Guerinot, M. L., Briat, J.-F., & Curie, C. (2002). IRT1, an Arabidopsis Transporter Essential for Iron Uptake from the Soil and for Plant Growth. The Plant Cell, 14(6), 1223–1233. 10.1105/tpc.001388

Wang, H. Y., Klatte, M., Jakoby, M., Bäumlein, H., Weisshaar, B., & Bauer, P. (2007). Iron deficiency-mediated stress regulation of four subgroup Ib BHLH genes in Arabidopsis thaliana. Planta, 226(4), 897–908. 10.1007/s00425-007-0535-x

Wang, W., Luo, L., Shi, H., Song, Y., Wang, J., Chen, C., Shen, Z., Rouached, H., & Zheng, L. (2024). The transcription factor OsSPL9 endows rice with copper deficiency resilience. Journal of Experimental Botany, 75(18), 5909–5922. 10.1093/jxb/erae273

Wang, Y. (2015). MIR156 Participates in Iron Homeostasis Through Targeting SPL9/SPL15 in Arabidopsis Thaliana.

Waters, B. M., & Armbrust, L. C. (2013). Optimal copper supply is required for normal plant iron deficiency responses. Plant Signaling & Behavior, 8(12), e26611. 10.4161/psb.26611

Waters, B. M., & Grusak, M. A. (2008a). Whole-plant mineral partitioning throughout the life cycle in Arabidopsis thaliana ecotypes Columbia, Landsberg erecta, Cape Verde Islands, and the mutant line ysl1ysl3. *New Phytologist*, *177*(2), 389–405. 10.1111/j.1469-8137.2007.02288.x

Waters, B. M., & Grusak, M. A. (2008b). Whole-plant mineral partitioning throughout the life cycle in Arabidopsis thaliana ecotypes Columbia, Landsberg erecta, Cape Verde Islands, and the mutant line ysl1ysl3. talic>New Phytologist, 177(2), 389–405. 10.1111/J.1469-8137.2007.02288.x

Waters, B. M., Uauy, C., Dubcovsky, J., & Grusak, M. A. (2009). Wheat (Triticum aestivum) NAM proteins regulate the translocation of iron, zinc, and nitrogen compounds from vegetative tissues to grain. Journal of Experimental Botany, 60(15), 4263–4274. 10.1093/jxb/erp257

Wu, G., Park, M. Y., Conway, S. R., Wang, J.-W., Weigel, D., & Poethig, R. S. (2009). The Sequential Action of miR156 and miR172 Regulates Developmental Timing in *Arabidopsis*. Cell, 138(4), 750–759. 10.1016/j.cell.2009.06.031

Wunderling, A., Ripper, D., Barra-Jimenez, A., Mahn, S., Sajak, K., Targem, M. B., & Ragni, L. (2018). A molecular framework to study periderm formation in Arabidopsis. New Phytologist, 219(1), 216–229. 10.1111/nph.15128

Xu, M., Hu, T., Zhao, J., Park, M.-Y., Earley, K. W., Wu, G., Yang, L., & Poethig, R. S. (2016). Developmental Functions of miR156-Regulated SQUAMOSA PROMOTER BINDING PROTEIN-LIKE (SPL) Genes in Arabidopsis thaliana. PLOS Genetics, 12(8), e1006263. 10.1371/journal.pgen.1006263

Zhai, Z., Gayomba, S. R., Jung, H.-I., Vimalakumari, N. K., Piñeros, M., Craft, E., Rutzke, M. A., Danku, J., Lahner, B., Punshon, T., Guerinot, M. L., Salt, D. E., Kochian, L. V., & Vatamaniuk, O. K. (2014). OPT3 Is a Phloem-Specific Iron Transporter That Is Essential for Systemic Iron Signaling and Redistribution of Iron and Cadmium in Arabidopsis. The Plant Cell. 10.1105/tpc.114.123737

Zhang, J., Liu, B., Li, M., Feng, D., Jin, H., Wang, P., Liu, J., Xiong, F., Wang, J., & Wang, H.-B. (2015). The bHLH Transcription Factor bHLH104 Interacts with IAA-LEUCINE RESISTANT3 and Modulates Iron Homeostasis in Arabidopsis. The Plant Cell, 27(3), 787–805. 10.1105/tpc.114.132704

